# Identification and characterization of a proliferative cell population in estrogen receptor-positive metastatic breast cancer through spatial and single-cell transcriptomics

**DOI:** 10.1101/2023.01.31.526403

**Authors:** Ryohei Yoshitake, Hitomi Mori, Desiree Ha, Xiwei Wu, Jinhui Wang, Xiaoqiang Wang, Kohei Saeki, Gregory Chang, Hyun Jeong Shim, Yin Chan, Shiuan Chen

## Abstract

**Background:** Intratumor heterogeneity is a hallmark of most solid tumors, including breast cancers. We applied spatial transcriptomics and single-cell RNA-sequencing technologies to profile spatially resolved cell populations within estrogen receptor-positive (ER^+^) metastatic breast cancers and elucidate their importance in estrogen-dependent tumor growth.

**Methods:** Spatial transcriptomics and single-cell RNA-sequencing were performed on two patient-derived xenografts (PDXs) of “ER-high” metastatic breast cancers with opposite estrogen-mediated growth responses: estrogen-suppressed GS3 (80–100% ER) and estrogen-stimulated SC31 (30–75% ER) models. The analyses included samples treated with and without 17β-estradiol. The findings were validated via scRNA-seq analyses on “ER-low” estrogen-accelerating PDX, GS1 (5% ER). The results from our spatial and single-cell analyses were further supported by the analysis of a publicly available single cell dataset and a protein-based dual immunohistochemical (IHC) evaluation using three important clinical markers [i.e., ER, progesterone receptor (PR), and Ki67]. The translational implication of these results was assessed by clinical outcome analyses on public breast cancer cohorts.

**Results:** Our novel space-gene-function study revealed a “proliferative” cell population in addition to three major spatially distinct compartments within ER^+^ metastatic breast cancers. These compartments showed functional diversity (i.e., estrogen-responsive, proliferative, hypoxia-induced, and inflammation-related). The “proliferative (*MKI67*^+^)” population, not “estrogen-responsive” compartment, was crucial for estrogen-dependent tumor growth, leading to the acquisition of luminal B features. The cells with induction of typical estrogen-responsive genes such as *PGR* were not directly linked to estrogen-dependent proliferation. Additionally, the dual IHC analyses demonstrated the distinct contribution of the Ki67^+^ proliferative cells toward estrogen-mediated growth and their response to palbociclib, a CDK4/6 inhibitor. The gene signatures developed from the proliferative, hypoxia-induced, and inflammation-related compartments were significantly correlated with worse clinical outcomes, while patients with the high estrogen-responsive scores showed better prognosis, confirming that the estrogen-responsive compartment would not be directly associated with estrogen-dependent tumor progression.

**Conclusions:** For the first time, our study elucidated a “proliferative” cell population distinctly distributed in ER^+^ metastatic breast cancers. They contribute differently toward progression of these cancers, and the gene signature in the “proliferative” compartment is an important determinant of luminal cancer subtypes.

## Background

Breast cancers are classified into at least four intrinsic subtypes by bulk gene expression profiling: luminal A, luminal B, HER2-enriched, and basal-like [1]. Each subtype presents distinct molecular features, clinical behaviors, and therapy responses. The luminal A and B estrogen receptor-positive (ER^+^) subtypes are distinguished by either relatively high or low estrogen-response gene expressions and low or high proliferative capacity, respectively. In clinical practice, a surrogate classification of breast cancer is widely performed using immunohistochemistry (IHC) for ER, progesterone receptor (PR; an estrogen-response marker), Ki67 (a proliferative marker), and HER2 status, serving as important criteria for therapy decisions [2].

ER^+^ breast cancers usually depend on estrogen for their development and progression [3]. Although the clinical outcomes of ER^+^ breast cancers have greatly improved by endocrine therapy, some patients, particularly those with metastasis, do not respond well to such treatments [4]. Furthermore, molecular basis of endocrine responses in “ER-low” (< 10% ER positivity) compared to “ER-high” tumors has not been clearly defined. Patient-derived xenografts (PDXs) with multiple cell types, including those from metastatic sites, are recognized as valuable for evaluating tumor response to a given agent, in addition to underlying mechanisms of action. We previously established a set of ER^+^ breast cancer PDXs with different levels of ER positivity and varying responses to estrogen [5,6]. These models enable a deeper understanding of the biology of ER^+^ breast cancers as well as the mechanisms of estrogen-dependent tumor growth, leading to better targeted treatment strategies.

Since current treatment approaches mostly target cancer as a homogeneous disease (e.g., endocrine therapy for ER^+^ breast cancers), intratumor heterogeneity often contributes to the acquisition of therapy resistance through the expansion of pre-existing resistant cells [7]. We reported that ER^+^ breast cancer PDXs contained both *ESR1*^+^ (encoding ER) and *ESR1*^-^ cells with molecular and functional differences using single-cell RNA sequencing (scRNA-seq) [8]. Nonetheless, the influence of estrogen on individual cell populations, as well as the importance of their responses toward the estrogen-dependent growth of ER^+^ breast cancers, has yet to be better defined.

Spatial transcriptomics (ST) is a promising approach which comprehensively and spatially characterizes the diverse cell populations on a given tissue section [9]. Visium, a widely used ST method from 10x Genomics, analyzes the transcriptome per each arrayed spot, which typically contains multiple (approximately 1–20) cells [10]. In contrast, scRNA-seq technology enables gene profiling at a single-cell level but does not preserve the spatial information of cells in tumor tissues due to the tissue dissociation process during the single-cell library preparation. Also, this dissociation process might introduce artificial gene expression via stress responses [9,11,12]. In addition, the translational value of scRNA-seq analysis on clinical specimens can be significantly affected because of their uncontrolled quality. Therefore, combining both ST and scRNA-seq analyses will provide a comprehensive view of the intratumor heterogeneity while maintaining spatial arrangement of cancer cells.

In this study, we performed ST and scRNA-seq analyses on PDX models with distinct responses to estrogen. Our ST analyses identified heterogeneous cell populations that were localized in different compartments of the tumor tissues and were commonly found both with and without estrogen treatment. Importantly, we estimated the space-gene-function relationship of these spatially resolved cell populations within ER^+^ tumors. These cell populations were further defined by scRNA-seq. Investigations using additional scRNA-seq datasets and *in vivo* experiments with a CDK4/6 inhibitor validated our findings and characterized distinct roles and responses to estrogen of the proliferative cells in ER^+^ breast cancers. Finally, we explored public clinical datasets using the gene signatures from the spatially distributed cell populations, providing the translational implications of the distinct functional compartments in ER^+^ metastatic breast cancers.

## Methods

### ST experiments on GS3 and SC31

Two “ER-high” PDXs, which were previously established from metastatic lesions and possessed definitive and contrasting growth responses to estrogen, were used (Supplementary Table S1). GS3 (80–100% ER) was E2-suppressive and luminal B-like, while SC31 (30–75% ER) was E2-dependent and luminal A-like [5,6,8]. Tumor pieces of GS3 and SC31 were implanted into mammary fat pads of 8–10-week-old NOD-SCID/IL2Rγ^−/−^ (NSG) mice, respectively [5,6,8]. Once the implanted tumors were established, placebo or 17β-estradiol (E2; 1 mg) pellet were implanted subcutaneously. To ensure that enough tumor mass remained for the analysis, GS3 tumors were implanted in intact mice and treated with E2 *in vivo* only for 7 days, whereas SC31 tumors were implanted in ovariectomized mice and treated for 6 weeks, as described in our previous scRNA-seq study [8]. After the treatment periods, the mice were euthanized, and tumor samples were collected.

Tumor tissues were trimmed to fit the fiducial frame of Visium Spatial Gene Expression Slide (10x Genomics, PN-2000233) while avoiding tumor core regions with large necrotic tissues. The tissue pieces were embedded on optimal cutting temperature compounds and stored at −80 °C. Cryosections were prepared at 10 μm thickness, stained with hematoxylin and eosin (H&E), and captured with Zeiss Observer II (Carl Zeiss). Tissue permeabilization was performed for 6 min as optimized using Visium Spatial Tissue Optimization Kit (10x Genomics, PN-3000394). The sequencing libraries from each section were prepared with Visium Spatial Gene Expression Reagent kit (10x Genomics, PN-1000187) according to the manufacturer’s instruction and sequenced with NovaSeq 6000 (Illumina). Raw sequencing data were processed using the 10x Genomics Space Ranger pipeline and Loupe browser and aligned to GRCh38 human or mm10 mouse genome.

### ST data analyses

ST data was processed using Seurat R package unless otherwise noted [13]. Details for initial assessment of the ST datasets are described in Supplementary Methods. The log-normalized data was obtained from the raw count data using NormalizeData function and used for the cell cycle analysis (using CellCycleScoring function) as well as visualization of gene expressions.

The spots in the ST dataset were classified based on *ESR1, PGR*, and *MKI67* expression (EPK classification). The spots without any count of *ESR1*, *PGR*, or *MKI67* genes in the raw count matrices were considered as “**N**egative (N)”, whereas the ones with any count (i.e., the raw counts ≥ 1) were considered as “**P**ositive (P)”. Each character of the class represents the status of *ESR1, PGR*, and *MKI67* expression (e.g., PNP represents a *ESR1*-positive, *PGR*-negative, *MKI67*-positive spot).

Raw count data on each tissue was normalized with SCTransform (SCT) function. The normalized data from the four tissues (GS3-Placebo, GS3-E2, SC31-Placebo, and SC31-E2) were integrated according to the developer’s vignette. Then, principal component analysis (PCA), Uniform Manifold Approximation and Projection (UMAP) dimension reduction, and cluster detection with the Louvain algorithm were performed (dims = 30, res = 0.6). To identify the genes highly expressed in each cluster, the differentially expressed genes (DEGs) were analyzed using the Wilcoxon rank sum test implemented in FindAllMarkers function on the SCT normalized data (logfc.threshold = 0.25, min.pct = 0.1). The top 10 genes for each cluster were visualized using DoHeatmap function.

To identify the gene signatures enriched in each cluster, the gene signature scores were calculated using VISION R package [14] and “hallmark” gene sets from Molecular Signature Database [15]. The Z score for each gene signature among clusters was calculated and visualized using Complexheatmap R package [16]. To assess the effect of E2 treatment on the gene signatures, the gene signature scores were compared between the individual clusters from placebo- and E2-treated tissues in each model.

### scRNA-seq analyses

The datasets prepared from GS3-Placebo/E2 and SC31-Placebo/E2 in our previous study were analyzed [8]. The “ER-low” GS1 dataset was prepared as described in Supplementary Methods. The human ER^+^ breast cancer dataset was obtained from the Broad Institute Single Cell portal [17]. After data integration (Supplementary Methods), PCA, UMAP dimension reduction, and cluster detection, as well as cell cycle analysis, DEG detection, VISION gene signature scoring, and EPK classification, were performed as described in the ST analyses. To compare the clusters identified in the ST dataset (ST clusters) and the clusters in scRNA-seq dataset (SC clusters) from GS3 and SC31, Pearson’s correlation coefficients of the hallmark gene signature scores were calculated between each ST and SC cluster and visualized using corrplot R package. The absolute value of a correlation coefficient between 0 to 0.2, 0.2 to 0.4, 0.4 to 0.6, 0.6 to 0.8, or 0.8 to 1.0 were considered as “no”, “weak”, “moderate”, “strong”, or “very strong” correlation, respectively.

Using the VISION R package and mammary “Stem” gene set curated in our previous article [18], the gene signature scores of the cells in each SC cluster were calculated. For the analyses on normal epithelium, a dataset established in our previous paper [18], which includes distinct epithelial lineages and their progenitors [i.e., basal, mammary stem cells/basal-progenitor (MaSC/B-pro), luminal hormone-sensing (L-Hor), L-Hor progenitor (LH-pro), luminal alveolar (L-Alv), and L-Alv progenitor (LA-pro)], was obtained. The mouse orthologs of human *ESR1/PGR/MKI67* (i.e., *Esr1/Pgr/Mki67*) were used for the EPK classification as described in the ST analyses.

### *In vivo* evaluation of palbociclib treatment on SC31

After the SC31 tumors were established, E2 pellets (1 mg) were implanted into the mice. Then, the mice were randomized into either E2 group (n = 5) or E2 + palbociclib group (n = 4). Palbociclib (LC Laboratories) was dissolved in phosphate-buffered saline and administered at 50 mg/kg daily via oral gavage. After 4 weeks of treatment, the mice were euthanized, and tumor samples were collected. The tumor tissues were fixed with 10% neutral buffered formalin and were embedded in paraffin.

### Dual IHC for ER/Ki67 and Ki67/PR

The dual IHC (ER/Ki67 and Ki67/PR) was performed by the Pathology Solid Tumor Core at City of Hope using Ventana Discovery Ultra IHC Auto Stainer (Roche Diagnostics). Primary antibodies used for the immunostaining include human ERα rabbit monoclonal antibody (Roche Diagnostics, 790-4325), human PR rabbit monoclonal antibody (Roche Diagnostics, 790-4296), and human Ki67 rabbit monoclonal antibody (Roche Diagnostics, 790-4286). Images were captured with VENTANA iScan HT (Roche Diagnostics) and analyzed using Cell Detection and Cell Classification functions implemented in Qupath software [19]. Staining protocol details are described in Supplementary Methods.

### Clinical outcome analysis on public cohorts with ST signatures

Molecular Taxonomy of Breast Cancer International Consortium (METABRIC) dataset [20] was obtained from cBioPortal [21,22] using cBioPortalData R package [23]. The gene signature scores on microarray data from each patient were calculated with GSVA R package [24] using the top 20 genes from the clusters identified in the ST analyses (Supplementary Table S2). The signature scores were compared between the intrinsic subtypes recorded in the METABRIC dataset. Then, the patients were divided into three groups based on the calculated scores, and the patients in the top and bottom tertiles were assigned as high and low groups, respectively. The patients were further classified based on the combination of ST_0 and ST_2 scores into four groups. Overall survival of each group was visualized with the Kaplan-Meier method. Details for additional cohorts [The Cancer Genome Atlas (TCGA) [25] and GSE124647 [26]] are described in Supplementary Methods.

### Statistical analysis

Statistical analyses were performed using R software. Two-group comparison was tested by Student’s t test and multiple comparison was adjusted by Bonferroni correction. Multiple group comparison was performed by one-way ANOVA followed by *post hoc* Tukey-Kramer test. The difference of overall survival among the groups was tested by log rank method. *P* < 0.05 was considered statistically significant.

## Results

### Estrogen dependency of ER^+^ breast cancers

To understand how different ER^+^ breast cancers respond to estrogen, we prepared and characterized a set of ER^+^ PDXs (Supplementary Table S1). These PDXs had varying levels of ER positivity and responses to E2 including: E2-dependent (SC31 [6], GS4), -accelerating (SC1 [6], GS1, GS2), and -suppressive (GS3 [8]) tumors (Supplementary Fig. S1A). These results support the clinical observation that not all ER^+^ patients respond to estrogen and endocrine therapies in the same fashion [4], and there is no direct correlation between ER levels and estrogen responsiveness. This points out the importance of understanding the molecular and cellular mechanisms of estrogen activity in tumors/tumor models with different response patterns, which leads to more effective targeted therapies.

### Spatial transcriptomics on two ER^+^ metastatic breast cancer PDX models with opposite responses to estrogen

To identify phenotypical diversity within ER^+^ breast cancers and to reveal how E2 affects their growths, we performed ST analyses on “ER-high” GS3 and SC31, both treated with and without E2 (Fig. 1A). Although PDX models are known to have “phenotypic drift” [27], growth suppression of GS3 and promotion of SC31 by E2 were maintained in the passages used in this study (Supplementary Fig. S1B).

**Figure 1.**
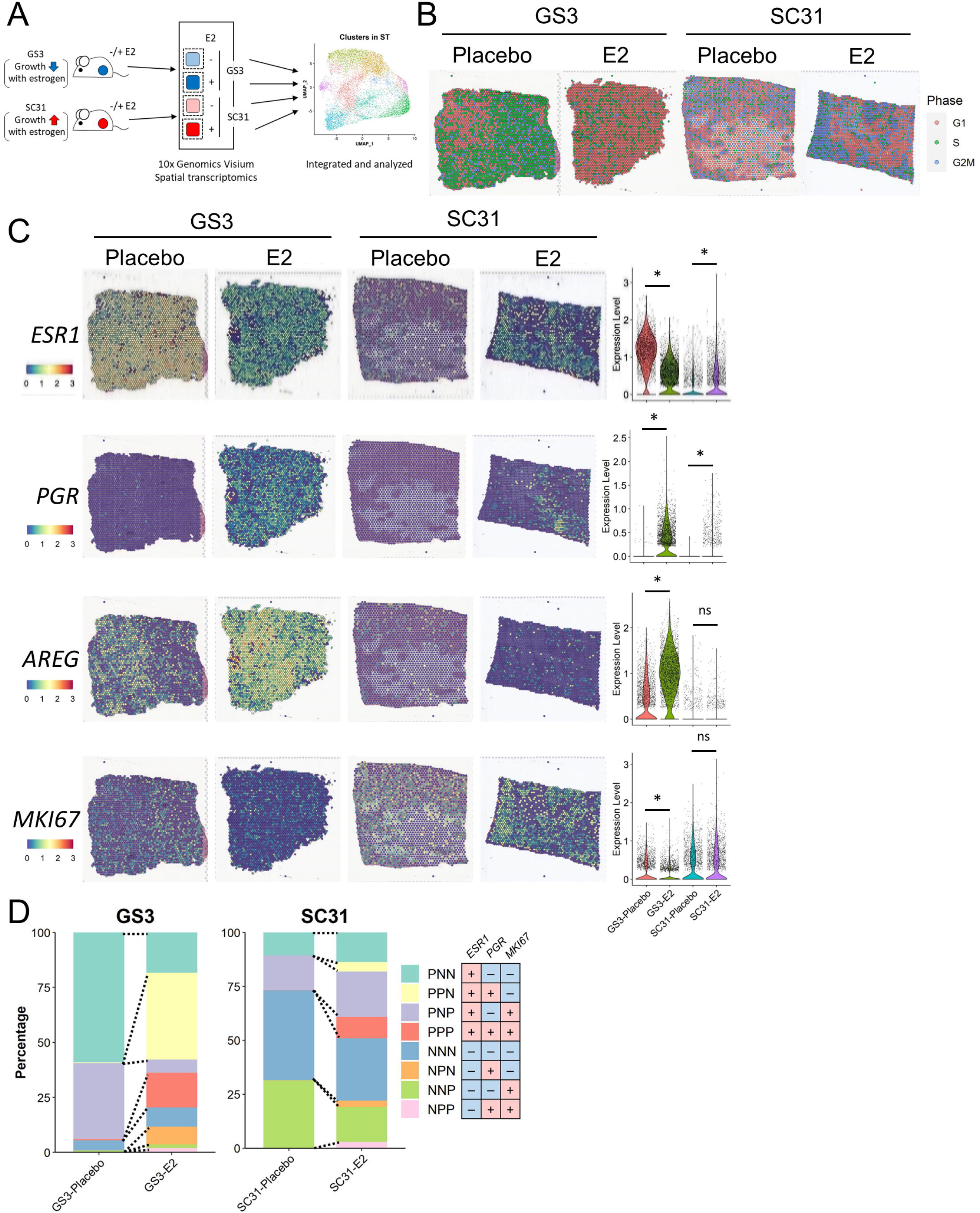
Spatial transcriptomics on two ER^+^ breast cancer PDX models. **A** The schematic workflow of our ST experiments. **B** Cell cycle analysis on the ST datasets from GS3 and SC31 with and without E2 treatment. The color of each spot represents the cell cycle phase. **C** Expression of *ESR1, PGR, AREG*, and *MKI67* genes in the ST datasets. The spatial plot on the left shows the location and level of the expression on each tissue. The violin plot on the right summarizes the gene expressions in each section. *, *P* < 0.05; ns, not significant. **D** EPK classification on the ST datasets.

To assess the quality of our ST experiments, we first evaluated the H&E images (Supplementary Fig. S2). GS3 sections mostly consisted of tumor cells and small patches of stromal and/or necrotic areas. Whereas in SC31 sections, we observed both tumor cells and scar-like areas which were possibly derived from the replacement of necrotic tissues during tumor growth. In the datasets obtained from the four tissue sections, a total of 9,323 spots were included (2,330 spots from GS3-Placebo, 2,863 spots from GS3-E2, 1,967 spots from SC31-Placebo, and 2,163 spots from SC31-E2) with an average number of 19,327 nCount and 4,824 nFeature for human genes. Although the spots located on the necrotic and/or scar-like areas showed lower values of both nCount and nFeature, the spots consisting of tumor cells were of better values in the quality control (QC) metrics (Supplementary Fig. S3), ensuring that the overall experimental procedures were successful.

During growth, PDX tumors are known to acquire mouse-derived stromal cells [27]. In our previous scRNA-seq datasets, single-cells isolated from GS3 and SC31 tumors included 10.6% and 16.2% mouse-derived stromal cells, respectively [8]. In the ST datasets, we observed the mouse-derived stromal cell marker expressions (Supplementary Fig. S4), but almost no human-derived stromal cell marker expressions (Supplementary Fig. S5). However, less than 5% of the spots included more mouse than human genes (Supplementary Fig. S6). In addition, these “mouse” spots were mostly located near the edge of tumor areas. These results demonstrated that the cells consisting of the ST spots in our datasets were mainly human-derived cancer cells and the impact of mouse stromal cells was minimal. Therefore, in this study, we focused on the analyses on the human transcriptome from the cancer cells.

### Spatial transcriptomics identified functional compartmentalization in ER^+^ metastatic breast cancers

To confirm the estrogen-dependent growth responses in the ST datasets, we performed a cell cycle analysis and examined the expression patterns of *ESR1, PGR*, and *MKI67* (encoding three important clinical markers ER, PR, and Ki67, respectively) (Fig. 1B and C). Our results indicated that the proportion of proliferating spots (S and G2M_phases) greatly decreased in GS3 and increased in SC31 with E2 treatment, confirming that our ST datasets reflected the discrete tumor growth responses of these PDX models (Fig. 1B and Supplementary Fig. S7). Furthermore, *MKI67* expression decreased in GS3 by E2 treatment and was maintained in SC31 (Fig. 1C). We found that the number of *ESR1*^+^ spots and its expression levels decreased in GS3 while increased in SC31 in E2-treated specimens, consistent with our previous scRNA-seq results [8]. Remarkably, *PGR* expression was found in both GS3 and SC31, but only when treated with E2. As a comparison, another well-known estrogen-regulated gene, *AREG*, was found in tumors without E2 treatment, especially in GS3, and was up-regulated by E2 treatment. Therefore, although known estrogen-regulated genes can be expressed in the absence of estrogen, the expression of *PGR* is completely dependent on estrogen. These observations are important to indicate that estrogen-regulated genes can be up-regulated independently from the overall tumor growth responses. *IL24*, an estrogen-induced suppressor as indicated in our previous report [8], was only detected in GS3-E2, supporting the conserved features of our PDX models (Supplementary Fig. S8). Next, we classified each spot by their *ESR1 /PGR/MKI67* gene expression patterns (Fig. 1D). Strikingly, the proportions of *PGR*^+^ spots, including *ESR1*^+^*PGR*^+^*MKI67*^-^ (PPN; each capital represents the positive/negative expression of each gene), PPP, NPN, and NPP spots, were increased by E2 treatment in both GS3 and SC31. While the other spot classifications (i.e., PNN, PNP, NNN, and NNP) could be identified in both placebo- and E2-treated tissues, only the changes in the proportion of PNP spots (i.e., a decrease in GS3 and an increase in SC31) correlated with the suppression of GS3 and promotion of SC31 by E2 treatment, respectively, indicating the potential importance of the cells in PNP spots (i.e., *ESR1*^+^ and/or *MKI67*^+^ cells) for estrogen-dependent tumor growth.

To reveal cell population(s) that drive estrogen-dependent breast cancer growth, we performed unsupervised clustering on the integrated dataset from the four ST samples (Fig. 1A). We identified nine clusters (ST_0–8) with diverse gene expression profiles (Fig. 2A and Supplementary Fig. S9A). Among the identified nine clusters, ST_3, ST_6, and ST_8 showed lower QC metric values (Supplementary Fig. S9B) and were mainly located in the necrotic and/or scar-like tissue areas (Fig. 2A and Supplementary Fig. S10), thus were not interrogated in further analyses. The other clusters (ST_0, ST_1, ST_2, ST_4, ST_5, and ST_7) were localized in the tumor areas, indicating that these clusters represented the heterogeneous populations of human breast cancer cells (Fig. 2A and Supplementary Fig. S10). Notably, each cluster was identified across the four samples and accumulated in a different compartment on the tumor sections (e.g., ST_0 spots on GS3-Placebo at the left area, whereas ST_2 at the right area). Particularly, ST_5 showed a unique localization surrounding the necrotic areas across the four samples. Importantly, our results showed intratumor compartmentalization of breast cancer cells which was shared irrespective of the different growth responses and presence of E2 treatment.

**Figure 2.**
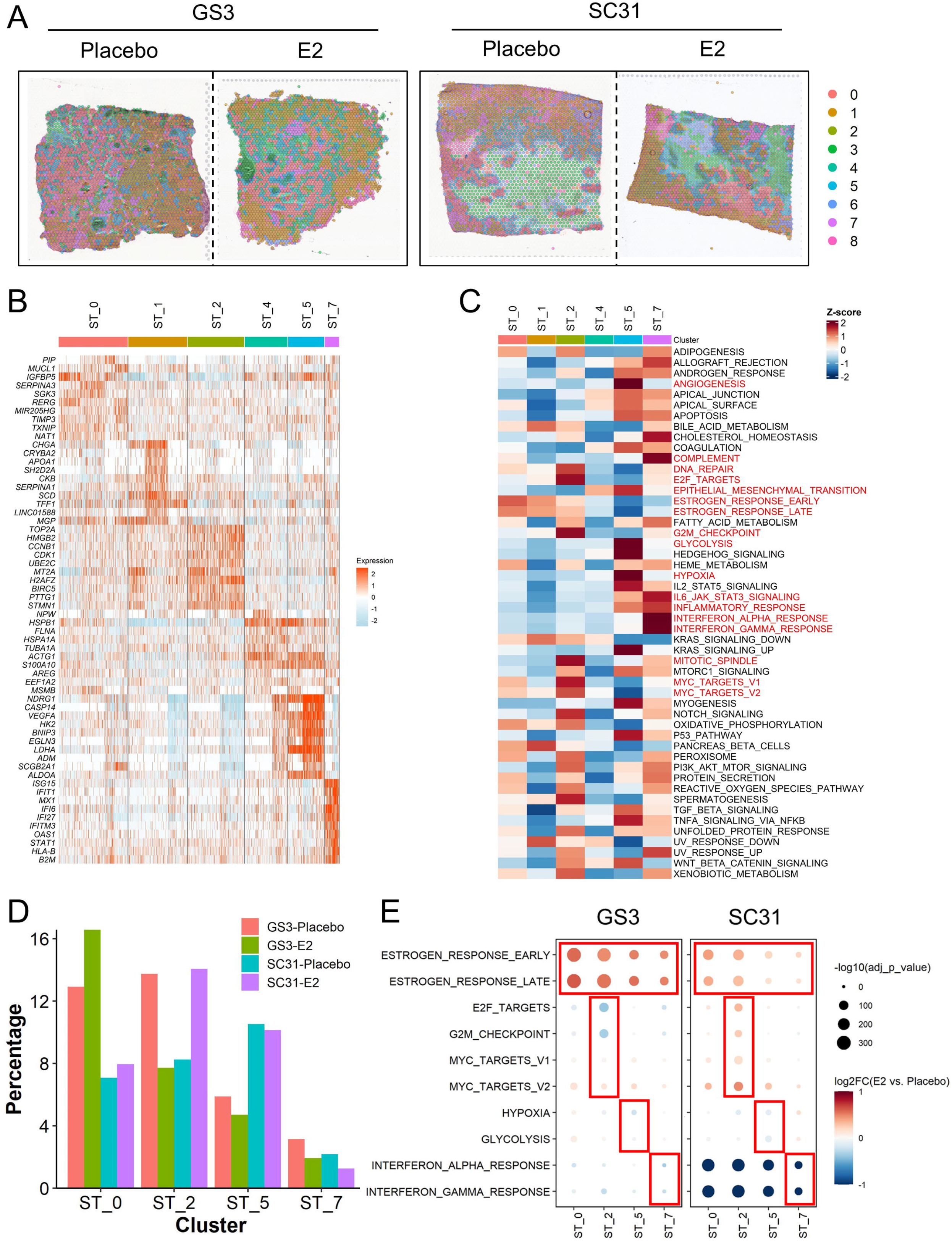
Identification of functional compartments in the ST dataset and the effect of E2 treatment. **A** Distribution of the ST clusters on tumor sections. Each spot is colored according to the cluster (ST_0–8) identified by the unbiased clustering. **B** Heatmap of the top 10 genes for the ST clusters. Column represents the spots and row represents the genes. The gene expression levels were scaled by SCTransform function. **C** Heatmap of the hallmark gene signature scores for the ST clusters. Column represents the ST clusters and row represents the gene sets. **D** Proportion of the four major ST clusters in each sample. The percentage of the ST cluster was calculated by (the number of spots in a ST cluster in a sample) / (the total number of spots in a sample). **E** Gene signature changes by E2 treatment. Size and color of the dot represent *P* value and log2 fold change value between E2 and placebo groups in each ST cluster, respectively.

To characterize the populations identified in our integrated dataset, we investigated the DEGs and gene signatures in each ST cluster (Fig. 2B and C). ST_0 showed higher expressions of *TXNIP* and *RERG*, which previous studies, including ours, reported as breast cancer suppressors associated with better prognosis [28–31]. Among the six main clusters, ST_0 also presented the highest scores for the estrogen-response gene sets (ESTROGEN_RESPONSE_EARLY/LATE), suggesting that ST_0 is the most responsive population to estrogen. The DEGs of ST_2 included many proliferation-related genes (e.g., *TOP2A, HMGB2*, and *CDK1*). Correspondingly, ST_2 had the highest scores in the gene signatures related to cell cycle progression (DNA_REPAIR, E2F_TARGET, G2M_CHECKPOINT, and MITOTIC_SPINDLE) as well as the highest percentage of the proliferating spots in the cell cycle analysis (Supplementary Fig. S11). They also showed the highest scores in MYC target signatures (MYC_TARGETS_V1/V2), which are known to be involved in breast cancer proliferation [32]. Additionally, ST_2 showed relatively higher scores for estrogen response gene sets among the six clusters, implicating their significance in the estrogen-dependent tumor proliferation.

ST_5 was characterized by higher expression of genes related to hypoxia and the resulting anaerobic cell metabolism or angiogenesis (e.g., *NDRG1, LDHA*, and *VEGFA*), as well as their associated gene signatures (ANGIOGENESIS, EPITHELIAL_MESENCHYMAL_TRANSITION, GLYCOLYSIS, and HYPOXIA). In previous studies, these features were reported to be connected to breast cancer metastasis [33,34]. Considering the localization of this cluster along with the necrotic/scar-like areas, ST_5 would represent the subset of cancer cells exposed to hypoxic microenvironments and thereby obtaining metastatic features. ST_7 showed higher gene expression of interferon (IFN)-inducible genes such as *ISG15, IFIT1*, and *IFI6*, which were recently demonstrated to be associated with ER^+^ breast cancer therapy resistance [35–37]. The ST_7 cluster also showed the highest scores in the signatures related to inflammation (e.g., COMPLEMENT, IL6_JAK_STAT3_SIGNALING, INFLAMMATORY_RESPONSE, and INTERFERON_ALPHA and GAMMA_RESPONSE). Although ST_1 showed higher expression of some estrogen-inducible genes (e.g., *TFF1*) and the estrogen-response gene signatures, their overall gene signature scores were lower than other clusters. The relatively lower scores were also observed in ST_4, implying that ST_1 and ST_4 would be functionally less distinct compared to the others. Collectively, the ST analyses revealed the four major spatially resolved “functional compartments” with important gene signatures identified in all four tumor sections.

### Impact of estrogen on individual functional compartments

Our ST analyses identified heterogeneous populations in either breast cancer models, regardless of estrogen treatment. Since E2 affected the growth of GS3 and SC31 in opposite ways, we compared the proportion of the four compartments (ST_0, ST_2, ST_5, and ST_7) between placebo- and E2-treated samples (Fig. 2D). ST_2 differed in accordance with the growth of the PDX models (i.e., decreased in GS3 and increased in SC31). In contrast, the proportions of ST_0, which was the compartment with the highest estrogen-response signature, was increased by E2 treatment in both models. Meanwhile, the abundance of ST_5 and ST_7 spots showed less changes by E2. Thus, our results indicated that the proliferative ST_2 population correlated with the E2-mediated tumor suppression of GS3 and promotion of SC31, respectively, and that the expansion of the estrogen-responsive ST_0 population was not directly linked to the tumor growth.

We then evaluated the effect of estrogen on gene expression profiles within each of the four major ST compartments by comparing the gene signature scores between the placebo- and E2-treated samples (Fig. 2E). In ST_2, the levels of genes associated with E2F_TARGETS and G2M_CHECKPOINT decreased in GS3 and increased in SC31 following E2 treatment. The levels of genes associated with ESTROGEN_RESPONSE_EARLY/LATE increased by E2 in both tumors. MYC_TARGETS_V1/V2 scores were highly increased in SC31 compared to that in GS3, suggesting the importance of these responses for inducing estrogen-dependent tumor proliferation. In ST_0, the levels of genes associated with ESTROGEN_RESPONSE_EARLY/LATE, but not the signatures related to proliferation, were increased by E2 in both models, supporting that the cells in ST_0 are not linked to E2-dependent tumor growth. The levels of genes associated with HYPOXIA and GLYCOLYSIS in ST_5 of both tumors were not much affected by E2, but ESTROGEN_RESPONSE_EARLY/LATE scores increased. While the major gene signature in ST_7 was INTERFERON_ALPHA and GAMMA_RESPONSE, E2 treatment decreased these signatures in all four ST clusters, especially in SC31. In summary, the scores of the estrogen response signatures increased across the ST clusters with E2, indicating that E2 treatment induced the typical estrogen-regulated gene expressions in most tumor cells in ER^+^ breast cancers, where both *ESR1*^+^ and *ESR1^-^* cells would be affected as reported in [8]. However, only the cells in ST_2 were associated with tumor growth. Furthermore, IFN-related gene signatures could be reduced by estrogen in many cells, which was also supported by the previous findings that the IFN-responsive genes were down-regulated by E2 in SC31 [8].

### Single-cell RNA sequencing defined the heterogeneous breast cancer population at a higher resolution

Recognizing the heterogeneous populations found in our ST dataset, we examined the scRNA-seq dataset integrated from the four tumor samples [8] and identified 14 clusters (SC_0-13) (Fig. 3A and Supplementary Fig. S12A). Despite the regression for stress response-related “immediate-early gene” (IEG) expressions (Supplementary Table S3 and Supplementary Methods), a small cluster with the high expression of IEGs (SC_8) persisted, and therefore, we did not interrogate this cluster to avoid potential artifacts in further downstream analyses (Supplementary Fig. S12B). Remarkably, the results of DEG analyses indicated that several SC clusters had comparable gene expression patterns with the ST clusters (Supplementary Fig. S12B and C); the top DEGs in ST_2 (*HMGB2*), ST_5 (*NDRG1*), and ST_7 (*ISG15*) were also highly expressed in SC_1/3/9, SC_10, and SC_13, respectively (Supplementary Fig. S12C). On the UMAP plot, there were three well-separated clusters, SC_1, SC_3, and SC_9, which showed highly proliferative signatures (Fig. 3A and B, and Supplementary Fig. S12B). Corresponding to the difference of their proliferative signature patterns (e.g., high DNA_REPAIR in SC_1 and high G2M_CHECKPOINT in SC_9, respectively), these clusters were further distinguished by their cell cycle phases: SC_1 in S phase, while SC_3 and SC_9 in G2M phase (Fig. 3B and C). SC_1 had the highest levels of estrogen response signatures among the three clusters, implying their different contribution toward estrogen-dependent growth.

**Figure 3.**
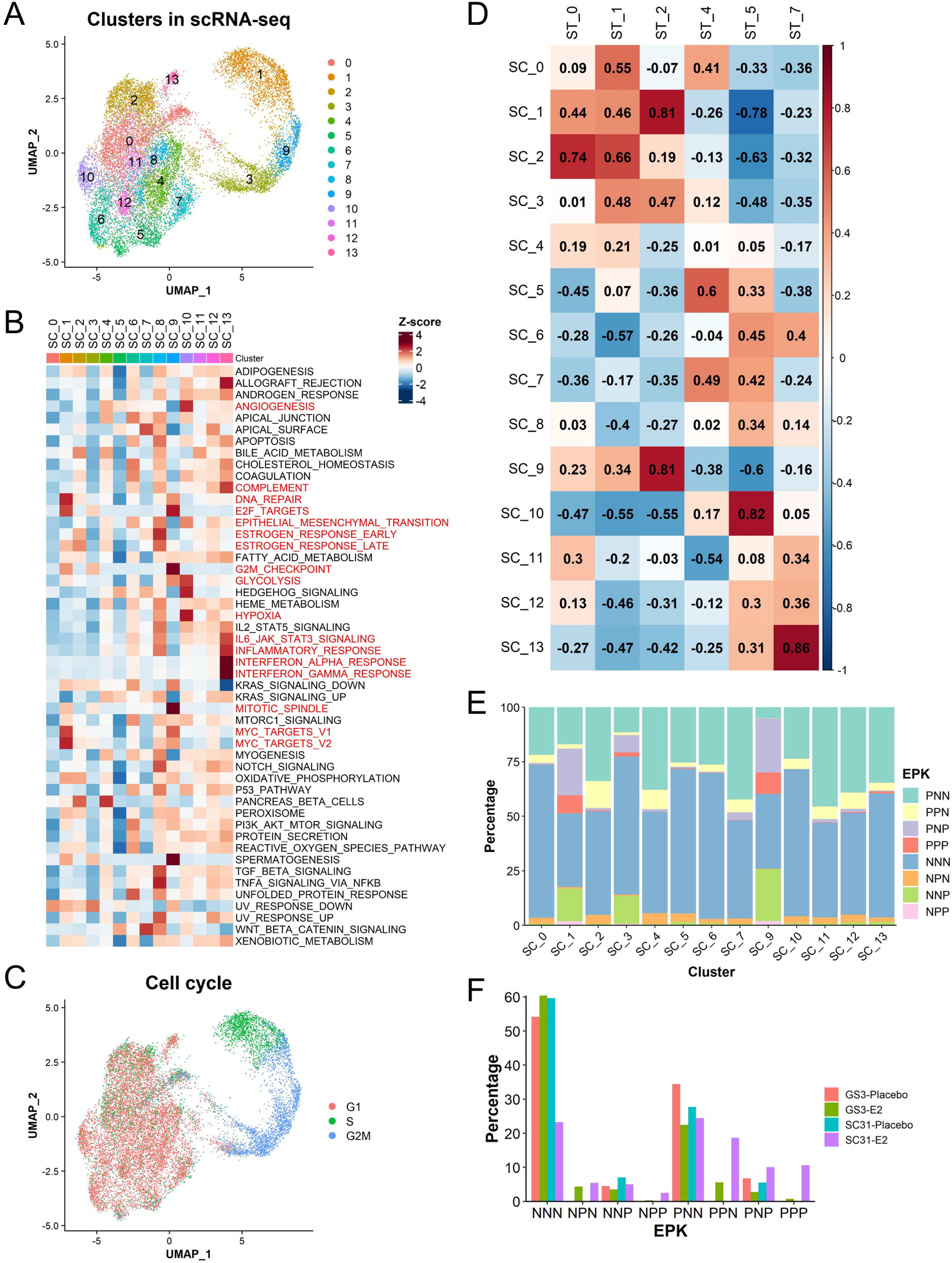
Single-cell analysis defined cell subsets responsible for estrogen-dependent tumor growth at a higher resolution. **A** UMAP plot of the integrated scRNA-seq dataset colored according to the clusters identified by the unbiased clustering. **B** Heatmap of the hallmark gene signature scores for the SC clusters. Column represents the SC clusters and row represents the gene sets. **C** Cell cycle analysis on the integrated scRNA-seq dataset. **D** Correlation analysis of the gene signature scores between the clusters in the ST and scRNA-seq datasets. The value and color of each box indicates Pearson’s correlation coefficient. **E** EPK classification on the scRNA-seq dataset. **F** Proportion of the cells with EPK classification in each scRNA-seq sample. The percentage of cells was calculated by (the number of cells in an EPK classification in a sample) / (the total number of cells in a sample).

To compare the clusters identified in the ST and scRNA-seq datasets, we analyzed the correlation of the gene signature scores between the ST and SC clusters (Fig. 3D). Corresponding to the similarity of the DEG and gene signature results, strong to very strong positive correlations were observed between ST_0 and SC_2 (estrogen-responsive), ST_2 and SC_1/9 (proliferative), ST_5 and SC_10 (hypoxia-induced), as well as ST_7 and SC_13 (inflammation-related). These correlations indicated the most dominant single-cell populations within the ST compartments. In addition, the ST clusters correlated with more than one SC cluster, indicating that the SC clusters defined cells included in each ST spot clusters at a higher resolution. Especially, ST_2 had very strong correlations with both SC_1 and SC_9 as well as a moderate correlation with SC_3, all of which depicted highly proliferative features. These results indicated that the ST_2 compartment would consist of three different proliferative cell subtypes. Altogether, we defined transcriptionally different compartments of breast cancer cells from the ST analysis and validated the concept that ST populations can be identified as distinct groups of cells, or their mixtures, using scRNA-seq datasets at a higher resolution.

### Further definition of cell subsets responsible for estrogen-dependent tumor growth with *ESR1 /PGR/MKI67* expressions

To better define “proliferative” ST_2-associated SC clusters vs. other clusters, we further investigated our scRNA-seq datasets using the classification based on *ESR1/PGR/MKI67* expressions. To evaluate the association of the expression of *ESR1 /PGR/MKI67* genes with the scRNA-seq clusters, we evaluated the distribution of cells in each EPK classification among the SC clusters (Fig. 3E). The *MKI67*^+^ cells (PNP, PPP, NPP and NNP) were specifically distributed among the proliferative SC clusters (SC_1, SC_3, and SC_9). The results indicated that the *MKI67*^+^ cells represented the proliferative populations essential for the estrogen-dependent tumor growth. In addition, SC_1 and SC_9 had more *ESR1^+^ /MKI67*^+^ cells (PNP and PPP), while SC_3 tended to have less, representing their differences in the estrogen responsive signatures. Meanwhile, *PGR*^+^ and *MKI67^-^* cells, including PPN and NPN, were present in all clusters except SC_1, SC_3, and SC_9, indicating that most *PGR*^+^ cells were not associated with proliferation.

To evaluate the effect of E2 on each group of EPK-classified cells, we analyzed the changes in their proportions by E2 treatment in our scRNA-seq dataset (Fig. 3F and Supplementary Fig. S13). PPN cells were only identified in E2-treated samples, ensuring that *PGR* expression was solely dependent on the presence of estrogen. NNP cells did not show major changes in their proportions with E2 treatment in both GS3 and SC31 models. Meanwhile, the changes in PNP cells correlated with the response of GS3 and SC31 to E2 treatment. PPP/NPP cells were mainly observed in SC31 (a luminal A-like subtype) with E2 treatment, also suggesting their roles in the proliferative response to estrogen. Accordingly, our results implicated that PNP, as well as PPP/NPP cells, are subsets of *MKI67*^+^ proliferative cells crucial for estrogen-dependent breast cancer growth.

### Validation of the results from SC31 and GS3 through single-cell analysis on another PDX and publicly available human breast cancer datasets

To validate what we have learned from the analyses on GS3 and SC31 PDXs, we performed scRNA-seq analysis on another PDX model named GS1. This “ER-low” (5% ER) GS1 showed luminal-B like features [5] and its growth was partly facilitated by E2 (but not E2-dependent), defining GS1 as an estrogen-accelerating luminal breast cancer model (Supplementary Table S1 and Supplementary Fig. S1). In the scRNA-seq dataset from GS1, the *MKI67*^+^ cells accumulated in the two clusters with S and/or G2M phase cells (clusters 2, 4) (Fig. 4A-C), similar to those of SC_1, SC_3, and SC_9 (in the GS3 and SC31 dataset) (Fig. 3E). There were a limited number of cells expressing *ESR1* and *PGR* in GS1. When comparing the EPK classification of this dataset (Fig. 4D), NNP cells were more dominant than PNP cells in the proliferative clusters and the number of NNP cells increased after E2 treatment (Fig. 4E and Supplementary Fig. S13), supporting that tumor growth in GS1 is driven mainly in an ER-independent manner, but accelerated in the presence of E2. Importantly, we identified the clusters with gene signatures related to hypoxia (cluster 5) and inflammatory responses (cluster 1), confirming the ST_5 and ST_7-like cells in this breast cancer (Fig. 4F).

**Figure 4.**
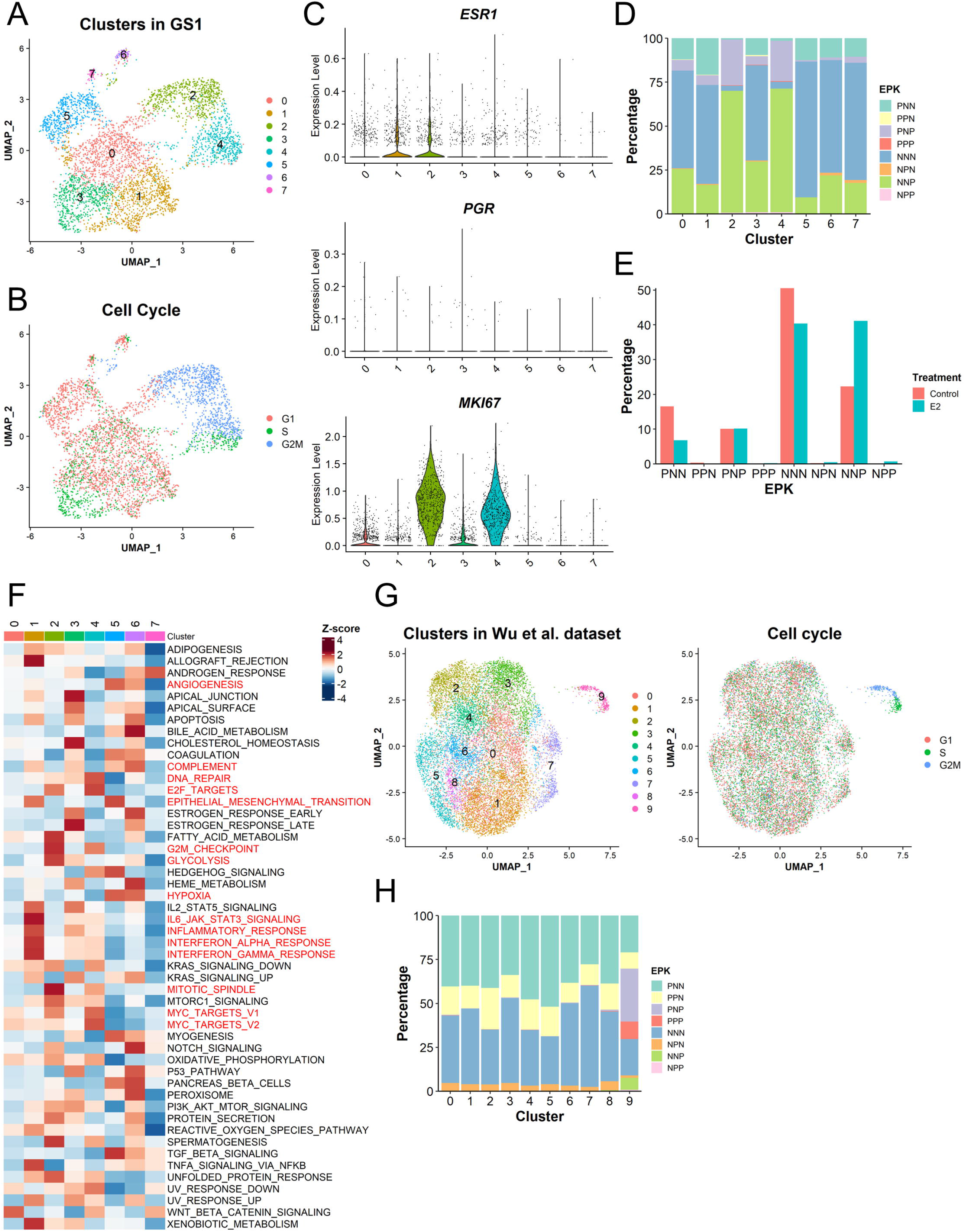
Validation of the results from SC31 and GS3 through single-cell analysis on two additional datasets. **A** UMAP plot of the GS1 dataset colored according to the clusters identified by the unbiased clustering and **B** the cell cycle phases. **C** Expression of *ESR1, PGR*, and *MKI67* expression in each cluster identified in the GS1 dataset. **D** EPK classification on the GS1 dataset. **E** Proportion of cells with EPK classification in each sample. The percentage of cells was calculated by (the number of the cells in an EPK classification in a sample) / (the total number of the cells in a sample). **F** Heatmap of the hallmark gene signature scores for the clusters in GS1. Column represents the clusters and row represents the gene sets. **G** UMAP plot of human ER^+^ breast cancer dataset colored according to the clusters identified by the unbiased clustering and the cell cycle phases. **H** EPK classification on the human ER^+^ breast cancer dataset.

The most abundant cell classifications in the entire datasets were NNN and PNN (Supplementary Fig. S13). These cells were present across all clusters, suggesting that these two cell classes are not functionally consistent, and that the genes specific to each cluster, other than *ESR1/PGR/MKI67*, would define their characteristics. Also, there were small fractions of NPN and NPP cells, which showed *PGR* expression despite the lack of *ESR1* expression. Although we need further validation of these cell classes, they would indicate the ER-independent mechanisms of estrogen for PR induction or the downregulation of ER expression after gaining PR.

To support our findings that the availability a function-specific “proliferative” cell population for the estrogen-dependent growth, we investigated an additional scRNA-seq dataset from nine ER^+^ breast cancer patient samples [17]. Among clusters identified in this dataset, cluster 9 showed the proliferative characteristics (Fig. 4G), although functional features of the other clusters were unclear due to varying sample quality and abundance of cancer cells (Supplementary Fig. S14A-C). EPK classification showed that the *MKI67*^+^ cells specifically accumulated in the proliferative cluster 9 (Fig. 4H), which confirmed the results from our own datasets (Fig. 3E). Importantly, *PGR*^+^ (e.g., PPN) cells were identified across all clusters regardless of different proliferative features, ensuring that *PGR* expression was not an indicator of proliferative capacity of the cells.

### Analysis of ER, PR, and Ki67 expression in SC31 through dual IHC following E2 and/or palbociclib treatment

To support the implication of *ESR1/PGR/MKI67* expressions at the protein level and evaluate the impact of targeted proliferative cell suppression, we performed an *in vivo* experiment using estrogen-stimulated SC31 treated with E2 and a CDK4/6 inhibitor, palbociclib (Fig. 5A and B, and Supplementary Fig. S15). The histological evaluation showed that the palbociclib treatment significantly reduced the total number of the cells per area (Fig. 5A and B), indicating that palbociclib successfully reduced tumor cell growth. With the palbociclib treatment, the dual ER/Ki67 IHC revealed that the number of both ER^+^Ki67^+^ and ER^-^Ki67^+^ cells were significantly reduced (Supplementary Fig. S15A). Furthermore, in Ki67/PR dual IHC, Ki67^+^PR^-^ cells, including both NNP and PNP cells, were remarkably decreased (Fig. 5B and Supplementary Fig. S15B). The number of Ki67^-^PR^+^ cells were not significantly affected by the treatment. Surprisingly, the number of Ki67^+^PR^+^ cells, including both NPP and PPP cells, was not significantly reduced by palbocilib, suggesting that palbociclib may not suppress these two classifications of cells, or that PR expression is linked to the non-proliferative features. Overall, these observations provided protein-based evidence for the ST/scRNA-seq results that the induction of the population with typical estrogen responses (e.g., *PGR*^+^ cells) was not directly linked to the estrogen-dependent growth and indicated that the therapeutic effect of palbociclib could be exerted by suppressing PNP and NNP cells.

**Figure 5.**
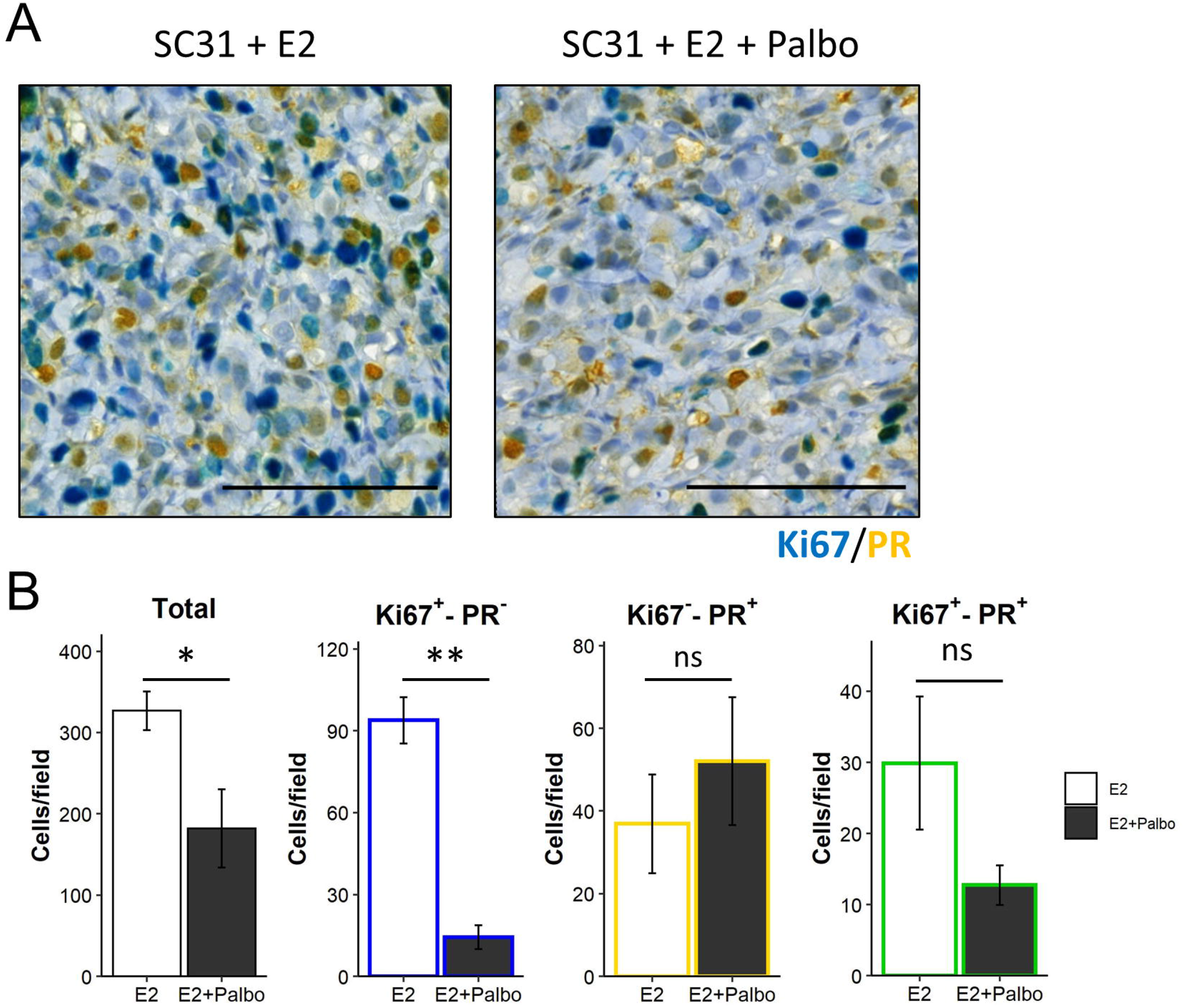
Dual IHC analysis on SC31 following E2 and/or palbociclib treatment. **A** Representative images of the dual IHC of Ki67 (blue) and PR (yellow) in SC31 treated with E2 or E2 + palbociclib (Palbo). The Ki67^+^PR^+^ cells were stained in green. Scale bar = 100 μm. **B** Quantification of the total and Ki67^+^PR^-^, Ki67^-^PR^+^, and Ki67^+^PR^+^ cells per field. Data are shown as mean ± SEM [n = 5 (E2) and 4 (E2 + Palbo)]. *, *P* < 0.05; **, *P* < 0.01; ns, not significant.

### Translational application of the four major functional compartments in ER^+^ breast cancers

To assess the clinical implication of our findings on the four functional compartments, we first analyzed a publicly available dataset, METABRIC, using the gene signatures from ST_0, ST_2, ST_5, and ST_7 compartments (Fig. 6A). Among the luminal breast cancer patients, the ST_0 signature scores were higher in the luminal A patients, while the scores of the other signatures (from ST_2, ST_5, and ST_7) were significantly higher in the luminal B patients. Particularly, the difference of the ST_2 (i.e., *MKI67*^+^) signature scores was highly significant, which agreed with the molecular definition of luminal B subtype as a highly proliferative ER^+^ breast cancer. The high score group of ST_2, ST_5, and ST_7 signatures had significantly shorter survival, suggesting these populations’ contributions toward the aggressive behaviors of ER^+^ breast cancers. However, patients with the high ST_0 scores showed better prognosis, confirming that ST_0 would not be directly associated with estrogen-dependent tumor progression.

**Figure 6.**
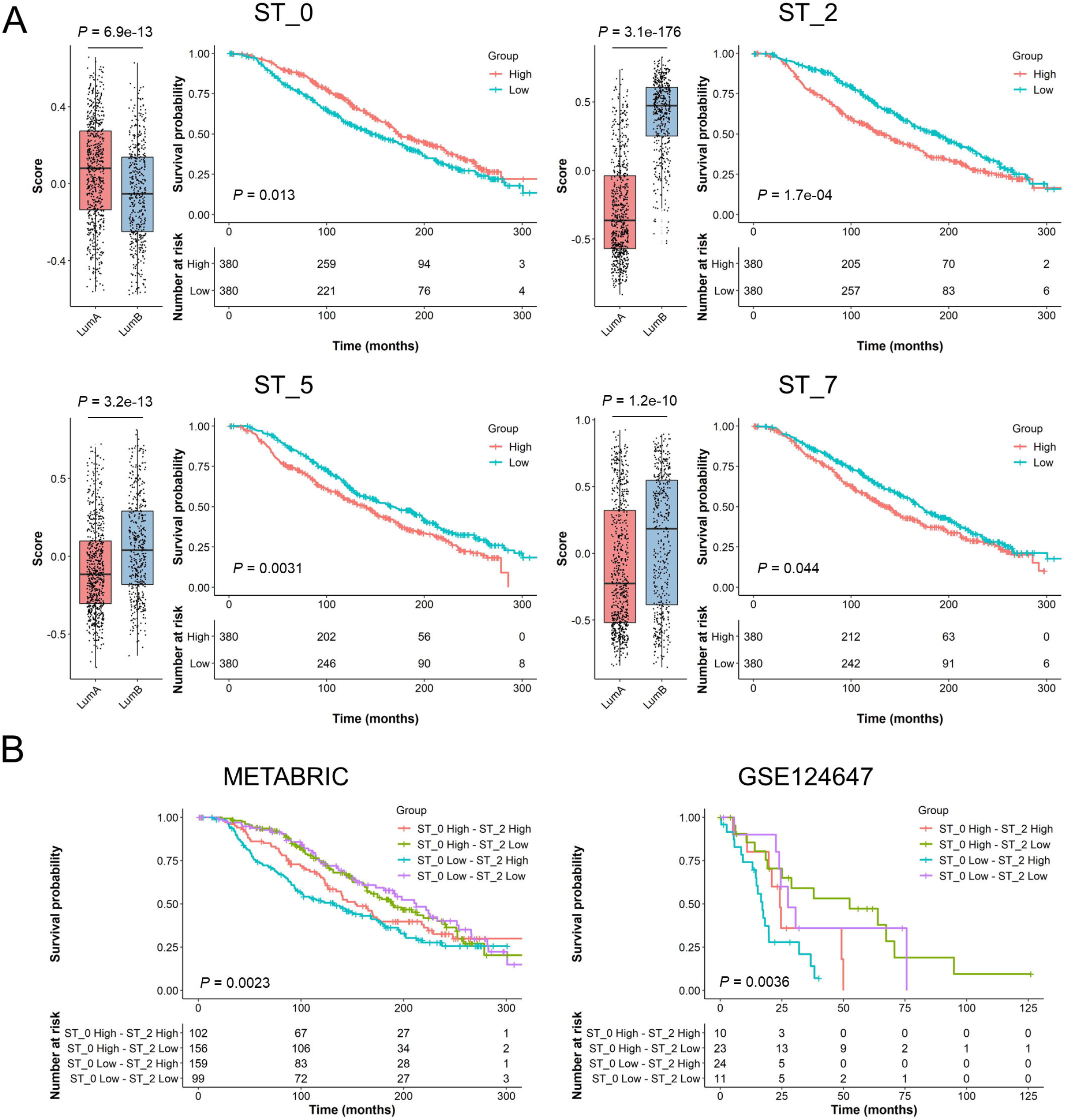
Clinical data analysis on public cohorts with the ST signatures. Luminal subtype breast cancers in METABRIC cohort (**A** and **B**; luminal A, n = 679; luminal B, n = 461) and stage IV ER^+^/HER2^-^ breast cancers in GSE124647 cohort (**B**; n = 140) were analyzed using **A** ST_0, ST_2, ST_5, or ST_7 signatures and **B** the combination of ST_0 and ST_2 signatures. The gene signature scores were calculated in each patient using GSVA R package. Left panels on **A** show the scores in luminal A and B patients. Kaplan-Meier plots show the overall survival of patients in each group.

Considering the importance of ST_0 and ST_2 signatures (i.e., estrogen response and proliferation) in luminal cancers and their opposite association with patient survival, we carried out an analysis combining these two gene signatures (Table 1 and Fig. 6B). The tumors with low ST_2 signature scores were primarily luminal A subtype with better prognosis, while the ST_0 signature scores did not show much impact. Furthermore, the tumors with high ST_2 scores, especially ST_0 low-ST_2 high tumors, were mainly luminal B subtype with shorter survival. We performed a similar comparison using TCGA data set (Table 1). The ST_2 low tumors were all luminal A subtype except for one tumor. ST_0 scores barely affected the luminal subtype distribution. We also examined a third dataset (GSE124647) that included 140 stage IV ER^+^/HER2^-^ breast cancer patients [26]. While this dataset did not have intrinsic subtype information, high ST_0 and ST_2 scores were associated with better and worse survival outcomes, respectively (Supplementary Fig. S16), the same as those observed in the METABRIC dataset (Fig. 6A). Similarly, ST_2 low tumors had a better outcome and ST_0 low-ST_2 high tumors had the shortest survival (Fig. 6B). The results from these analyses indicated that ST_2 (including *MKI67*^+^ cells) had more impact than ST_0 on luminal tumor prognoses, and luminal B tumors could be associated with a higher abundance of ST_2 compartment. Also, it was clear that luminal A tumors had low ST_2 scores.

**Table 1.** ST_0 and ST_2 combination groups and luminal subtypes in the public cohorts.

|  | METABRIC |  | TCGA |  |
| --- | --- | --- | --- | --- |
|  | Luminal A | Luminal B | Luminal A | Luminal B |
| ST_0 High - ST_2 High | 27<br>(26%) | 75<br>(74%) | 23<br>(42%) | 32<br>(58%) |
| ST_0 High - ST_2 Low | 152<br>(97%) | 4<br>(3%) | 103<br>(99%) | 1<br>(1%) |
| ST_0 Low - ST_2 High | 22<br>(14%) | 137<br>(86%) | 21<br>(22%) | 73<br>(78%) |
| ST_0 Low - ST_2 Low | 96<br>(97%) | 3<br>(3%) | 64<br>(100%) | 0<br>(0%) |

Our analysis of ER^+^ PDXs, including ST and scRNA-seq analyses, determined *MKI67*^+^ ST_2 compartment as the major driver for the proliferation of ER^+^ cancers. We previously established a normal mammary epithelial cell atlas and curated gene signatures representing distinct epithelial lineages [18]. Our analyses have suggested that luminal A cancers have their origins in the L-Hor cell lineage, and luminal B type is more associated with the progenitor state (LH-pro) [18]. Remarkably, in the current study, the proliferative SC clusters (SC_1, SC_3, and SC_9) had higher scores for the mammary “Stem” signature developed from the epithelial atlas (Fig. 7A), indicating the similarities between these *MKI67*^+^ cells and the normal mammary stem/progenitor-like cells. In the normal mammary epithelial cell atlas, we also found that the *Mki67*^+^ cells were only observed in the progenitor populations (Fig. 7B). NNP cells were found in all three progenitor populations, whereas PNP, NPP, and PPP cells were only found in the LA-pro and LH-pro populations. NPP and PPP cells were more abundant in the LH-pro population than in the LA-pro population, suggesting their more differentiated features toward L-Hor lineage.

**Figure 7.**
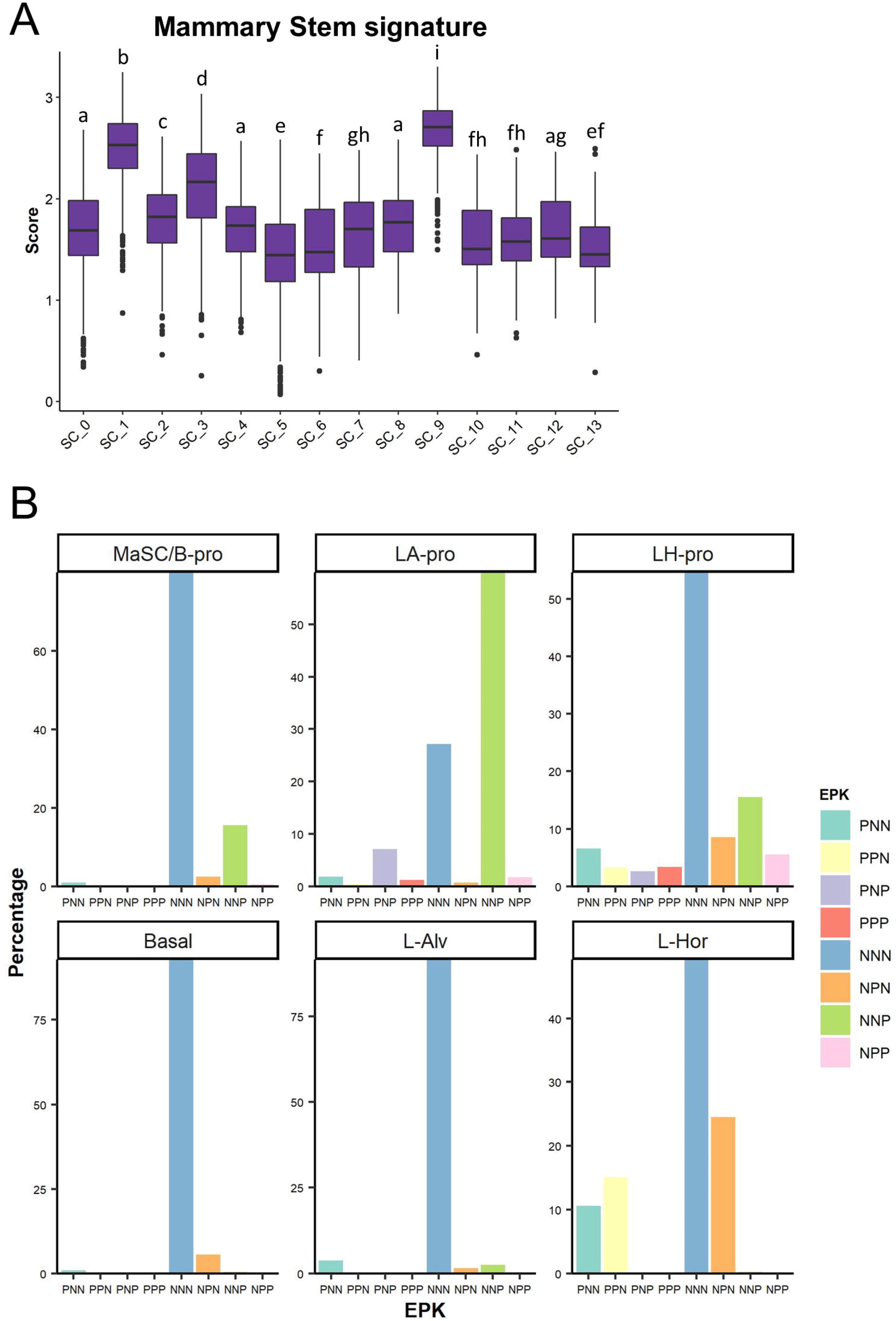
Progenitor-like features of the *MKI67*^+^ proliferative cell subsets. **A** Mammary “Stem” gene signature scoring on the GS3/SC31 scRNA-seq dataset. The gene list for the mammary stem signature was derived from [18] and the score was calculated using VISION R package. Different letters on the box plots indicates significant difference between the groups (*P* < 0.05). **B** EPK classification of normal mammary epithelium dataset. Each panel shows the results on each lineage of cells identified in our previous study [18].

Further investigation on the entire METABRIC cohort including other subtypes (i.e., basal-like, claudin-low, and HER2-enriched) showed that luminal A and B had higher scores in ST_0 signature than the other subtypes, and again, the clinical outcome of high score group was better than the low group’s (Supplementary Fig. S17). For ST_2 signature, luminal A was the lowest, and luminal B had relatively higher scores amongst the subtypes. Although ST_2 was characterized as an estrogen-responsive and proliferative population, this signature was also high in the basal-like subtype because the gene sets for ST_2 signature consisted of many proliferation-related genes. ST_5 score was significantly higher in the basal and HER2 tumors, which depicted their more aggressive and metastatic behaviors as well as estrogen-independent features. Also, the high ST_5 score indicated the worse outcome, supporting the potential contribution of ST_5 to cancer metastasis via estrogen-independent mechanisms. The difference of ST_7 signature scores among the subtypes were also significant but less clear than the other signatures. In addition, the levels of the ST_7 signature did not show significant value on the survival of all breast cancer patients. It may indicate that the activation of their inflammation-related features, especially IFN-responsive characters, was important only for the aggressiveness of luminal breast cancers. In summary, our analysis on the clinical cohorts supports the translational significance of the results from our ST analyses on the ER^+^ metastatic breast cancer PDXs.

## Discussion

In this study, we revealed the functional compartmentalization within ER^+^ metastatic breast cancer by the application of ST and scRNA-seq analyses on three unique models, including two high-ER^+^ PDXs (GS3 and SC31) with opposite responses to estrogen, followed by validation through an estrogen-accelerating low ER-expressing PDX (GS1). Using these patient tumor-derived models with definitive estrogen responses and containing both ER^+^ and ER^-^ cells, we identified four spatially separated compartments (i.e., estrogen-responsive, proliferative, hypoxia-induced, and inflammation-related), which showed unique gene signatures as previously reported mechanisms of cancer development and progression [32–37]. Among our important findings, the responsiveness of a highly proliferative cell population (i.e., ST_2) to estrogen was shown to be crucial for ER^+^ cancer growth, leading to the acquisition of luminal B features. The “proliferative” cell population was further validated using an independent published single cell dataset [17]. Through the analysis of clinical databases, the four major compartments demonstrated their contributions toward predictive patient outcomes in a distinct manner, proving the translational importance of our preclinical results.

Clinically, ER^+^ breast cancer is typically defined as 1% or higher ER expression, and 10% ER expression is considered the threshold for endocrine therapy response. While our study has confirmed ER-positivity indicates a dominant proliferative population in a tumor, co-expression of ER and Ki67 is essential for estrogen-dependent proliferation. Surprisingly, the cells with the induction of well-known estrogen-regulated genes (e.g., *PGR*) were not directly associated with the estrogen-dependent growth. Also, the protein-based evaluations suggested that PNP and NNP cells could be linked to CDK4/6 inhibitor therapy responses in ER^+^ breast cancers.

The association between the estrogen-regulated gene expression (e.g., *PGR, AREG*) and the estrogen-dependent growth has been extensively investigated [38–42]. Previous studies mainly used breast cancer cell lines, especially MCF-7, which are known to show concomitant upregulation of estrogen-responsive genes and cell proliferation genes within the same cell responding to estrogen. The results in the current study on tumor models with heterogenous gene expression indicated that the expansion of cells expressing typical estrogen-regulated genes like *PGR* would not be directly associated with estrogen-mediated proliferation. Expression of estrogen-regulated genes could indicate a better response to the endocrine therapy, as we observed a better clinical outcome associated with ST_0 gene signature (Fig. 6). Our findings support the observation in clinical practice that luminal A (low ST_2 gene signature) patients with higher estrogen-regulated gene expressions (e.g., *PGR*) show better prognosis compared to luminal B (high ST_2 gene signature) patients with low estrogen response and high proliferation [3]. While Scabia et al. revealed the **inter**-patient tumor heterogeneity of the hormone response [43], our ST and scRNA-seq data revealed the contribution of function-associated **intra**-tumor heterogeneity for patient-specific hormone responses. Although progesterone was indicated to suppress the estrogen-dependent proliferation of MCF-7 cells [44], a progesterone treatment did not affect the proliferation nor the bulk gene expression pattern of estrogen-suppressive GS3 (Supplementary Fig. S18A and B). Overall, our results highlight the power of ST technology for identifying physically separated functional compartments in the translational models.

The *MKI67*^+^ cells were exclusively included in the SC clusters correlating with ST_2. Our results supported the previous clinical study that only Ki67, but not ER/PR, was the negative factor for patient prognosis in IHC3, an IHC-based scoring method [45]. Among the *MKI67*^+^ cells, we identified two distinct classifications of the cells correlated with estrogen-dependent growth, i.e., PNP and PPP/NPP cells. The scRNA-seq analyses indicated that PNP cells could be observed both with and without E2, and they were implied to be associated with estrogen-dependent tumor growth in ER^+^ breast cancers. Meanwhile, PPP/NPP cells were suggested to be solely estrogen-dependent and only present in a less aggressive luminal A-like subtype. Intriguingly, our *in vivo* experiment showed that palbociclib could suppress PNP, but not PPP/NPP cells, providing the potential explanation why palbociclib has been only effective for advanced breast cancers and highlight the importance of its combination with endocrine therapy to target both PNP and PPP/NNP populations in ER^+^ breast cancers. The additional examination on the estrogen-accelerating, but not totally dependent, GS1 indicated that NNP cells could be the dominant cell type in driving cancer growth in “ER-low” breast cancers. It is noted that the NNP cell population in GS1 increased after estrogen treatment as it implied potential interactions between *ESR1*^+^ and *ESR1*^-^ proliferative cells. In the normal mammary epithelium dataset, NNP cells were found in the mammary stem or basal progenitor populations, implying that they are present not only in luminal cancers but also in other subtypes such as basal-like cancers. Collectively, our study would indicate the complexity of cell distribution driving the tumor proliferation in ER^+^ breast cancers, i.e., ER-dependent (PNP and PPP/NPP) and -independent (NNP) cells, and our findings support the importance of combination therapy to suppress all proliferative populations within a single tumor.

Both GS3 and SC31 were derived from metastasized breast tumors. Our ST analysis found a compartment with higher hypoxia-induced signatures (i.e., ST_5). Hypoxia is one of the hallmarks of cancers and often occurs alongside necrosis formed by insufficient oxygen and nutrition supply. The hypoxia-induced phenotypical changes of breast cancer cells have been thought to be important especially for distal metastasis [33]. Indeed, our hypoxia-induced cluster concomitantly showed the enrichment of gene signatures known to promote metastasis. Notably, the hypoxia-related gene signature in ST_5 was not greatly affected by estrogen treatment in this study. Also, the non-luminal breast cancers in the METABRIC cohort showed higher expressions of the ST_5 gene signature, indicating the importance of hypoxic features as estrogen-independent mechanisms of tumor progression for all breast cancer subtypes. Our ST analysis has revealed that glycolysis is linked to ST_5, while oxidative phosphorylation is associated with proliferative ST_2 that requires efficient energy production (Fig. 2C). These results pose the opportunity for future combination treatment of endocrine therapy with hypoxia-targeting agents (e.g., anti-angiogenic therapy) to suppress metastasis.

Our ST analysis identified another compartment showing the inflammation-related signatures (i.e., ST_7), especially with the upregulation of IFN-responsive genes. IFNs have been investigated as inflammatory mediators produced by immune cells, facilitating the anti-tumor immunity. However, emerging evidence suggests that the tumor cells themselves can produce IFNs and thereby activate IFN signaling autonomously [46]. A recent study demonstrated that the potential autonomous activation of IFN signaling in ER^+^ breast cancers led to the CDK4/6 inhibitor resistance [36]. Also, the long-term inhibition of estrogen signaling in breast cancer cells can induce IFN-responsive gene expressions, leading to endocrine resistance [35,47]. In our study, we observed the downregulation of IFN-response signatures in ST_7 with E2 treatment. With considering that our SC31 model was partly suppressed by endocrine therapy in our previous study [6], we speculate that the IFN-responsive signaling would be usually suppressed by estrogen in ER^+^ breast cancers, but once the estrogen signaling was inhibited, the IFN-responsive population would become dominant for the breast cancer’s progression. While further mechanistic studies are needed, this study provides another layer of the therapeutic target toward improved clinical outcomes in ER^+^ breast cancers.

A limitation of this study is that we mainly focused on two ER^+^ breast cancer PDXs, GS3 and SC31, with confirmative evidence from another “low-ER” PDX GS1. The design of this study required well-defined estrogen-responsive models. Yet, consistent intratumor heterogeneity and functional compartmentalization was clearly demonstrated in these structurally and functionally distinct tumor models. While the translational value of scRNA-seq analysis on clinical specimens can be significantly affected because of their uncontrolled quality, the recent paper by Wu et al. performed large-scale scRNA-seq analyses on clinical samples and determined seven cell groups with distinct gene signatures called gene module (GM) across all breast cancer subtypes [17]. Importantly, their GMs corresponded to our ST/SC signatures (e.g., the proliferation-related gene signatures in GM3 or the IFN-responsive genes in GM4). Our analysis on the nine ER^+^ specimens from their dataset confirmed the presence of the “proliferative” cell population (i.e., cluster 9 in Fig. 4G), well-separated from other cell populations which were impacted by the heterogeneous number of cells from different patients.

Our comprehensive ST analyses on ER^+^ PDXs with defined estrogen responses, for the first time, revealed functionally distinct compartments within ER^+^ breast tumors and demonstrated the importance of the ST_2 signature, including Ki67 expression, using the METABRIC and two other cohorts. Again, the strength of our study using PDXs is the ability to evaluate the response to experimental intervention, i.e., estrogen treatment. These approaches allowed us to elucidate the essential cell populations for estrogen-dependent tumor growth. Such information is currently not available. While multiple important analyses on correlations between bulk gene profiling and clinical outcome have been carried out using primary tumor specimens, a limitation is the lack of direct evidence for gene expression-tumor growth relationship. Thus, our findings are crucial not only for improving the understanding of ER^+^ breast cancer biology, but also for providing insights to develop more targeted therapeutic strategies toward better patient prognoses.

## Conclusions

Our study identified the four active compartments in ER^+^ metastatic breast cancers (i.e., estrogen responsive, proliferative, hypoxia-induced, and inflammation-related) (Fig. 8). These four spatially separated functional compartments are suggested to uniquely contribute to the breast cancer progression. We identified a cell cluster linked to estrogen-dependent tumor proliferation that were not directly associated with the induction of typical estrogen-responsive genes. Moreover, the proliferative cells were further distinguished into more specific cell subsets based on the expression of *ESR1, PGR*, and *MKI67*, implicating their distinct contribution toward tumor growth. Overall, our study on estrogen-responsive PDXs through ST and scRNA-seq analyses provides a comprehensive view of intratumor heterogeneity in ER^+^ breast cancers and reveals the key functional compartments as important candidates for refining current treatment strategies.

**Figure 8.**
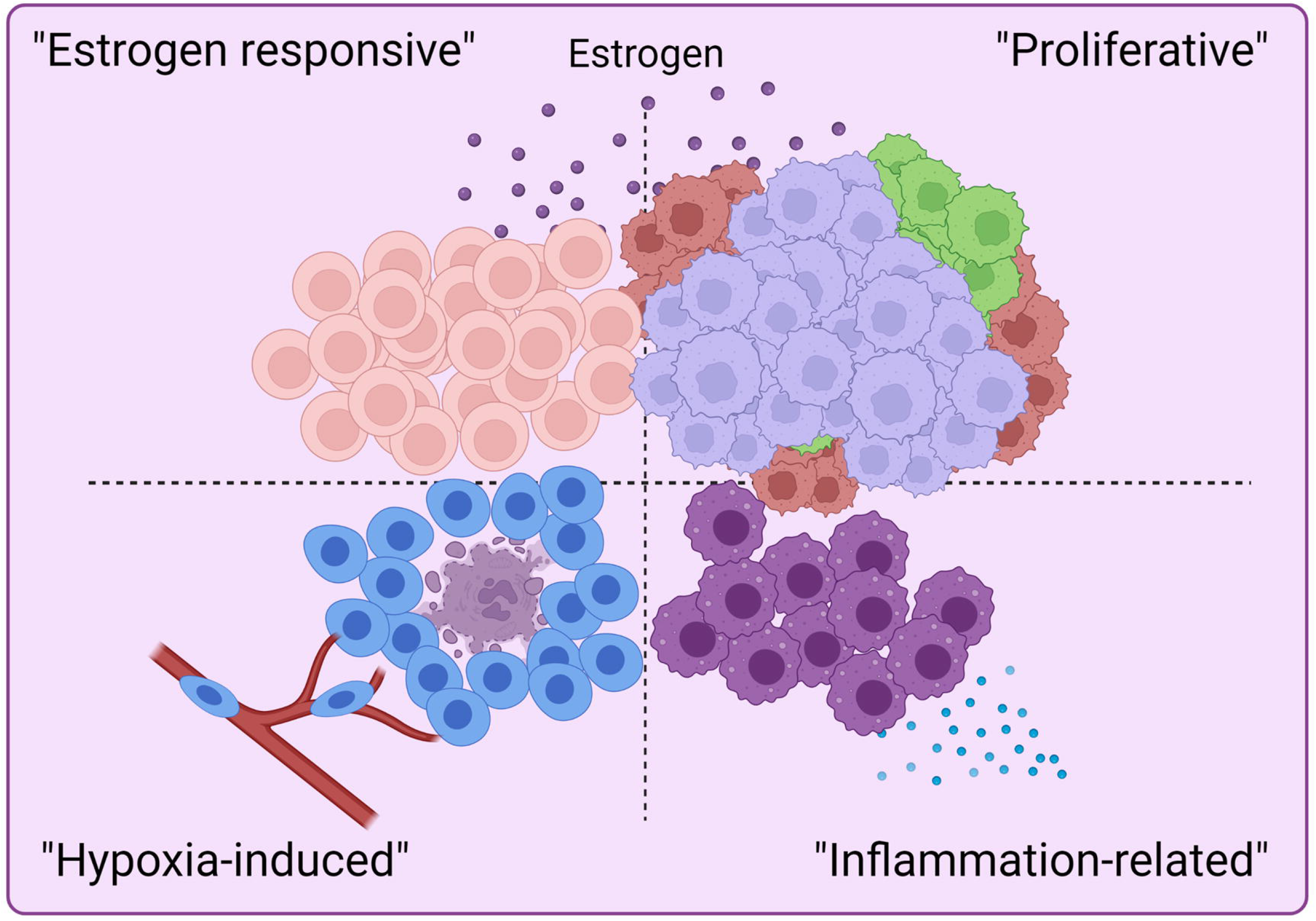
Graphical abstract. We identified four spatially separated compartments in ER^+^ breast cancers with distinct gene signatures, leading to overall cancer progression in a function-specific manner.

## Abbreviations

ST: Spatial transcriptomics
scRNA-seq: Single-cell RNA-sequencing
ER: Estrogen receptor
PDX: Patient-derived xenograft
IHC: Immunohistochemistry
PR: Progesterone receptor
E2: 17β-estradiol
NSG: NOD-SCID/IL2Rγ^−/−^
H&E: Hematoxylin and eosin
SCT: SCTransform
PCA: Principal component analysis
UMAP: Uniform Manifold Approximation and Projection
DEG: Differentially expressed gene
MaSC/B-pro: Mammary stem cells/basal-progenitor
L-Hor: Luminal hormone-sensing
LH-pro: Luminal hormone-sensing progenitor
L-Alv: Luminal alveolar
LA-pro: Luminal alveolar progenitor
METABRIC: Molecular Taxonomy of Breast Cancer International Consortium
TCGA: The Cancer Genome Atlas
QC: Quality control
IFN: Interferon
IEG: Immediate-early gene
GM: Gene module

## Data Availability

The ST datasets obtained in this study (GS3-Placebo, GS3-E2, SC31-Placebo, and SC31-E2) and the scRNA-seq datasets from GS1 have been deposited in the National Center for Biotechnology Information Gene Expression Omnibus data repository (GSE214571 and GSE213733, respectively).

## Authors’ contribution

RY and SC designed the study. RY, HM, HJS, and YC performed animal experiments. DH and X. Wang performed histological analyses. X. Wu and JW performed the sequencing. RY, X. Wu, KS, GC performed bioinformatic analyses. SC supervised the study. RY, HM, DH, X. Wang, KS, GC, YC, and SC drafted and revised the manuscript. All authors read and approved the final manuscript.

## Acknowledgments

This work was supported by The Lester M. and Irene C. Finkelstein endowment and NIH U01ES026137 (S. Chen). We are grateful to the City of Hope Core Facilities (supported by NIH P30CA033572), including the Integrative Genomics Core, Pathology: Solid Tumor Core, and Light Microscopy Digital Imaging Core, for the excellent technical support. We also thank Kiana Kelii and Justin Cheng for assistance with data analyses. Graphical abstract was created with BioRender.com.

