## Supplementary Methods for "Identification and characterization of a proliferative cell population in estrogen receptor-positive metastatic breast cancer through spatial and single-cell transcriptomics"

Evaluation of the effect of estrogen treatment on estrogen receptor-positive (ER^+^) breast cancer patient-derived xenograft (PDX) models

The PDXs were prepared as described in [1] and the clinical information of the original tumors are provided in Table S1. To assess the effect of estrogen on tumor growth, female NOD-SCID/IL2Rγ^−/−^ (NSG) mice with established tumors were treated with 17β-estradiol (E2; 1 mg) pellets. For the GS1 and GS2 models, the ovariectomized NSG mice were implanted with tumor pieces and E2 (1mg) pellets. For the GS4 model, the intact NSG mice were implanted with tumor pieces and E2 (1mg) pellets. Control groups were either implanted with placebo pellets (GS1 and GS2) or did not receive treatment (GS4). Tumor volumes were recorded to generate growth curves.

Initial assessment of ST datasets

Quality control metrics of human gene information, including the number of unique molecular identifier counts (nCount), the number of unique genes (nFeature), and the mitochondrial gene percentage per spot, were evaluated.

Mouse gene expressions and percentage of mouse gene counts in each spot were analyzed in the entire ST datasets. Since mouse gene expressions were sparsely present across the tissues, only human gene information was used for the other analyses in this study.

Single-cell RNA sequencing (scRNA-seq) dataset preparation

*GS3 and SC31*

The datasets prepared from GS3-Placebo/E2 and SC31-Placebo/E2 in our previous study were analyzed [2]. After filtering out cells with mitochondrial gene % > 20% and/or nFeatures < 200, the four datasets were normalized with SCTransform (SCT) function and integrated according to the Seurat’s SCT vignette. In the initial results, there were two clusters with either high mitochondrial genes or immediate-early genes (IEGs) (Table S3), which were both considered as artifacts from the single-cell processing [3–5]. Therefore, these two factors were regressed out during the SCT normalization before the integration as described in [5].

*GS1*

The GS1 tumor pieces were implanted into 11–13-week-old ovariectomized NSG mice as described above. After E2 treatment for 16 weeks, the tumor tissues were collected and were processed for single-cell preparation as described in our previous study [2]. The tumor tissues were cut into 2 mm thick strips and were enzymatically digested with 1.5 mg/mL DNAse I (Millipore Sigma, 10104159001), 0.4 mg/mL collagenase IV (Worthington Biochemical Corporation, CLS-4), 5% fetal bovine serum, and 10 mM HEPES in Hank’s buffered salt solution for 40 mins. After being strained through a 70 μm cell strainer, the samples were treated with Ammonium-chloride-potassium lysis buffer to remove residual red blood cells. Dead cells were removed with Dead Cell Removal MicroBeads (Miltenyi Biotec, 130-090-101). After the cell viability were ensured to be > 75% using TC20 Automated Cell Counter (Bio-Rad), the samples were loaded onto the Chromium Controller (10x Genomics) targeting 2,000–5,000 cells per lane. The Chromium v3 single-cell 3’ RNA-seq reagent kit (10x Genomics, PN-1000092) was used to generate single-cell RNA-seq libraries according to the manufacturer’s protocol. The libraries were sequenced with the NovaSeq 6000 system (Illumina). Raw sequencing data were processed using the 10x Genomics Cell Ranger pipeline (version 3.1.0) and then aligned to GRCh38 human genome or mm10 mouse genome. The single-cell samples from two biological replicates from each group (control and E2) were combined and processed for scRNA-Seq. The barcodes containing mouse genes were removed from the analysis. After filtering out cells with mitochondrial gene % > 30% and/or nFeatures < 3,000 or > 9,000, the SCT normalization, regression for IEG and mitochondrial gene expressions, and data integration were performed as described in the GS3/SC31 datasets.

*Human ER^+^ breast cancers*

A human breast cancer scRNA-seq dataset was obtained from the Broad Institute Single Cell portal [6]. After subsetting only normal and cancer epithelial cells from ER^+^ specimen, the SCT normalization, regression for IEG and mitochondrial gene expressions, and data integration were performed as described in the GS3/SC31 datasets.

Dual immunohistochemistry (IHC) for ER/Ki67 and Ki67/PR

The dual IHC (ER/Ki67 and Ki67/PR) was performed by the Pathology Solid Tumor Core at City of Hope using Ventana Discovery Ultra IHC Auto Stainer (Roche Diagnostics). Heat-mediated antigen retrieval was performed using Cell Conditioning 1 buffer (pH 8.5; Roche Diagnostics, 950-124) and the sections were incubated with primary antibodies. Two target antigens (ER/Ki67 or Ki67/PR) were sequentially detected, and the heat inactivation was performed to prevent the cross-reactivity between each antigen detection. Primary antibodies used for the immunostaining include: human ERα rabbit monoclonal antibody (Roche Diagnostics, 790-4325), human PR rabbit monoclonal antibody (Roche Diagnostics, 790-4296), and human Ki67 rabbit monoclonal antibody (Roche Diagnostics, 790-4286). After the primary antibody incubation, the sections were incubated with DISCOVERY anti-Rabbit HQ (Roche Diagnostics, 760-4815) or DISCOVERY anti-Rabbit NP (Roche Diagnostics, 760-4817) as well as DISCOVERY anti-HQ-HRP (Roche Diagnostics, 760-4820) or DISCOVERY anti-NP-AP (Roche Diagnostics, 760-4827). Stains were visualized with DISCOVERY Yellow Kit (Roche Diagnostics, 760-239) or DISCOVERY Teal Kit (Roche Diagnostics, 760-247), followed by the counterstain with hematoxylin. Images were captured with VENTANA iScan HT (Roche Diagnostics) and analyzed using Cell Detection and Cell Classification functions implemented in Qupath software [7].

Clinical data analysis on The Cancer Genome Atlas (TCGA) and GSE124647 cohorts

TCGA breast cancer RNA-seq data [8] was obtained using cBioPortal [9,10] using cBioPortalData R package [11]. The gene signature scores on RNA-sequencing data from each patient were calculated with GSVA R package [12] using the top 20 genes from ST_0 and ST_2 compartments (Table S2). The patients were divided into three groups based on the calculated scores, and the patients in the top and bottom tertiles were assigned as high and low groups for each gene signature, respectively. The patients were further classified based on the combination of ST_0 and ST_2 scores into four groups. The luminal subtypes were compared among the groups.

Another metastatic ER^+^ and HER2^-^ breast cancer cohort data [13] was obtained from National Center for Biotechnology Information (NCBI) Gene Expression Omnibus (GEO) data repository (GSE124647). The gene signature scores on microarray data from each patient were calculated as described in the METABRIC analysis. For a gene annotated with multiple probes, a mean value was calculated to represent its expression level. The group assignments based on ST_0, ST_2, and their combination were performed as described above. Overall survivals of each group were visualized with the Kaplan-Meier method.

Progesterone (P4) treatment on GS3 model

The intact NSG mice were implanted with GS3 tumor pieces as described above. After the tumor mass was established, the mice were treated with E2 (1mg) and/or P4 (10mg) pellets. The control group did not receive treatment. After the treatment for 4 weeks, the tumor tissues were collected and were processed for bulk RNA-sequencing (RNA-seq) as described in our previous study [2]. The bulk RNA-seq data was obtained from NCBI GEO data repository (GSE156922).
