## Supplementary Figure S1-18 for "Identification and characterization of a proliferative cell population in estrogen receptor-positive metastatic breast cancer through spatial and single-cell transcriptomics"

### Slide 1
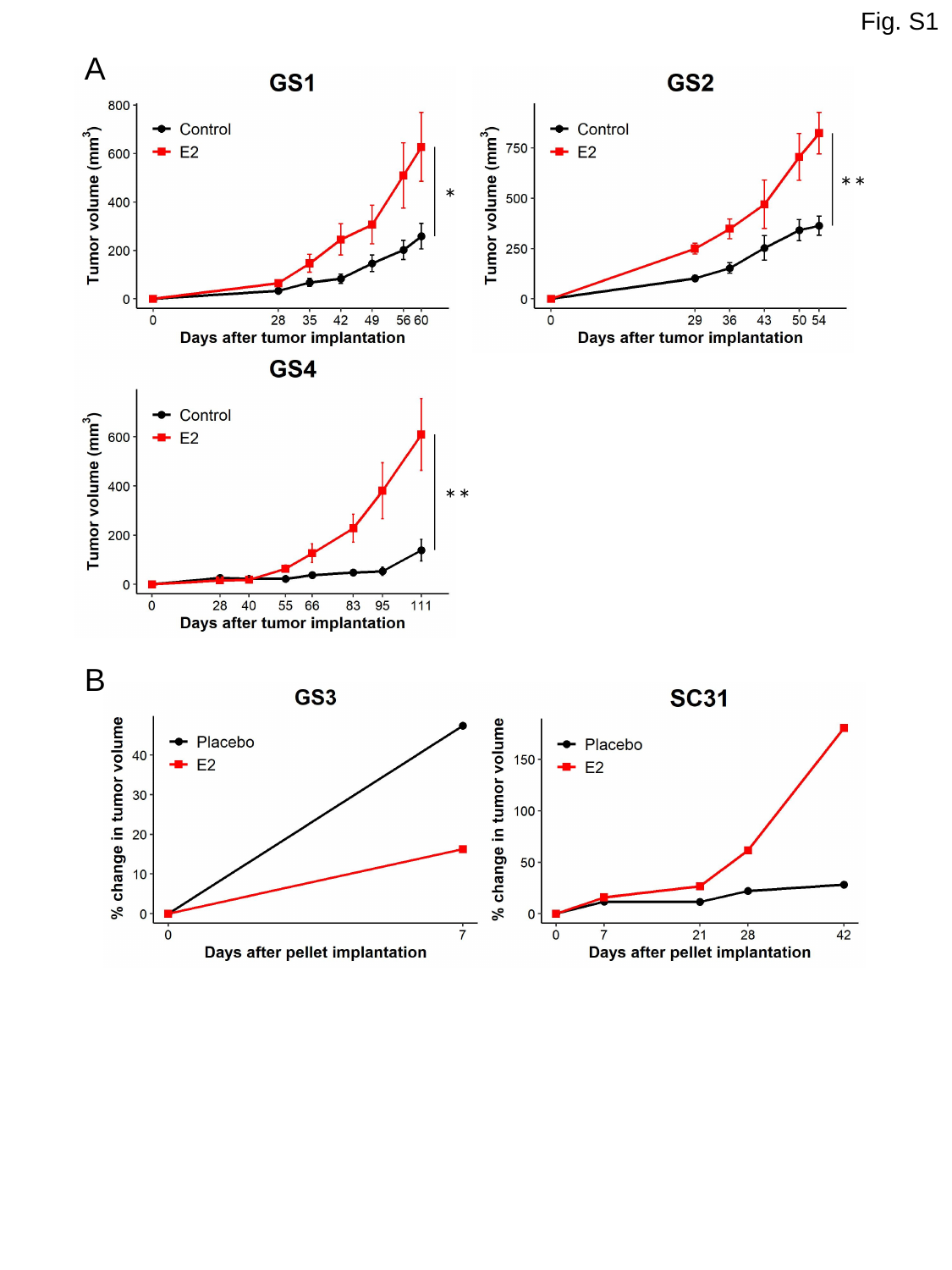

Fig. S1
A
**
*
**
B

### Slide 2
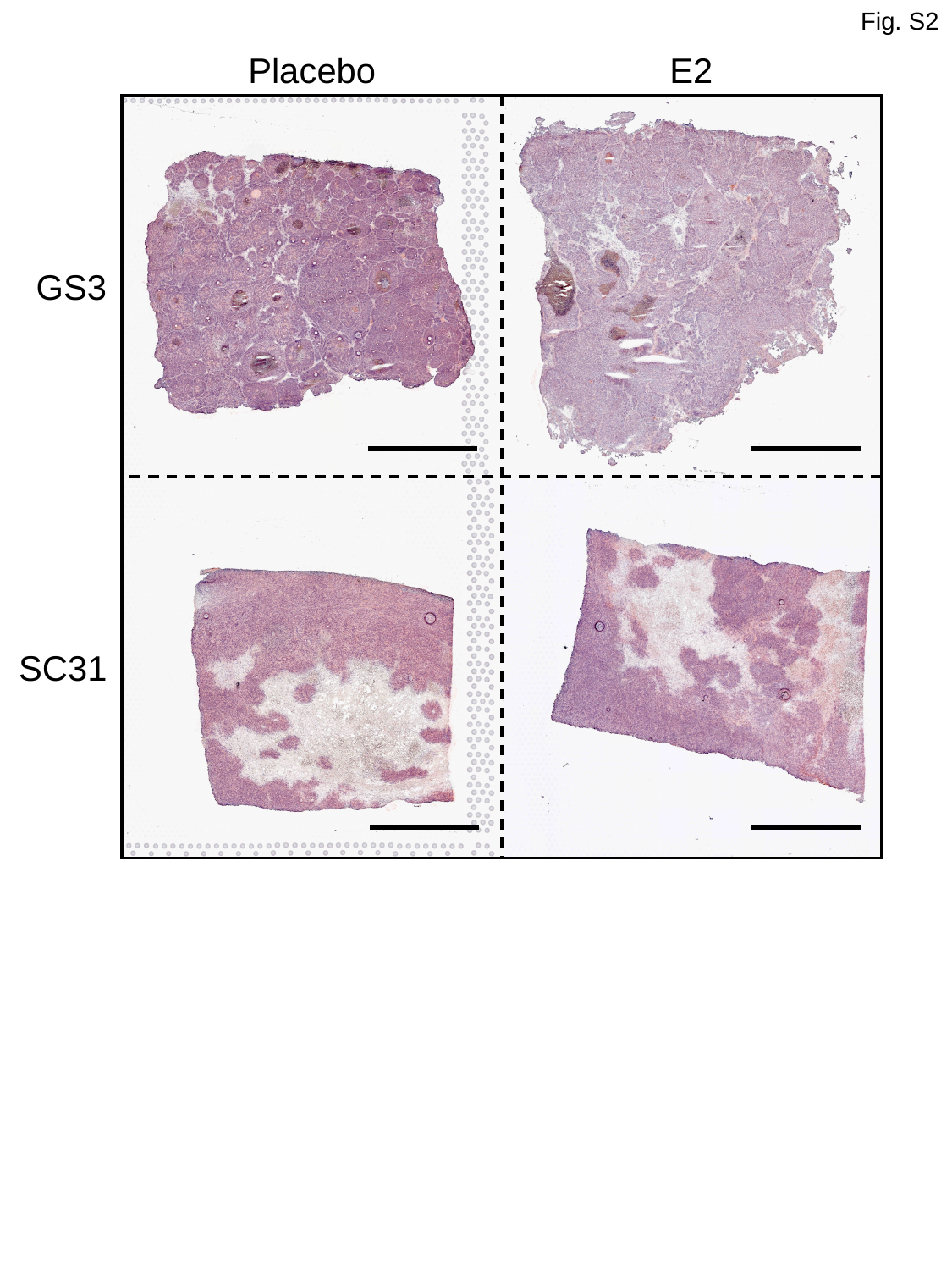

Fig. S2
Placebo
E2
GS3
SC31

### Slide 3
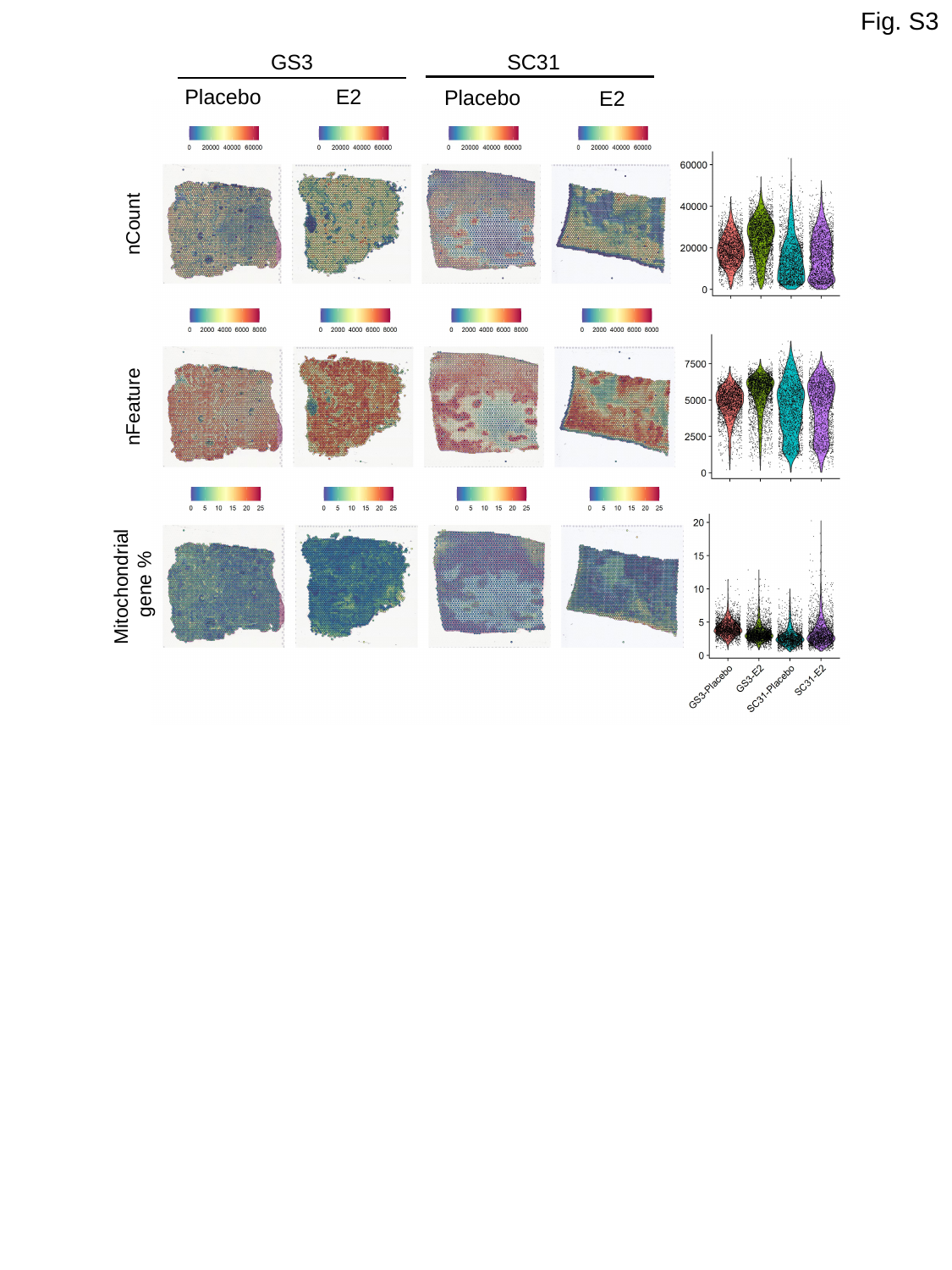

Fig. S3
GS3
SC31
E2
Placebo
Placebo
E2
nCount
nFeature
Mitochondrial
gene %

### Slide 4
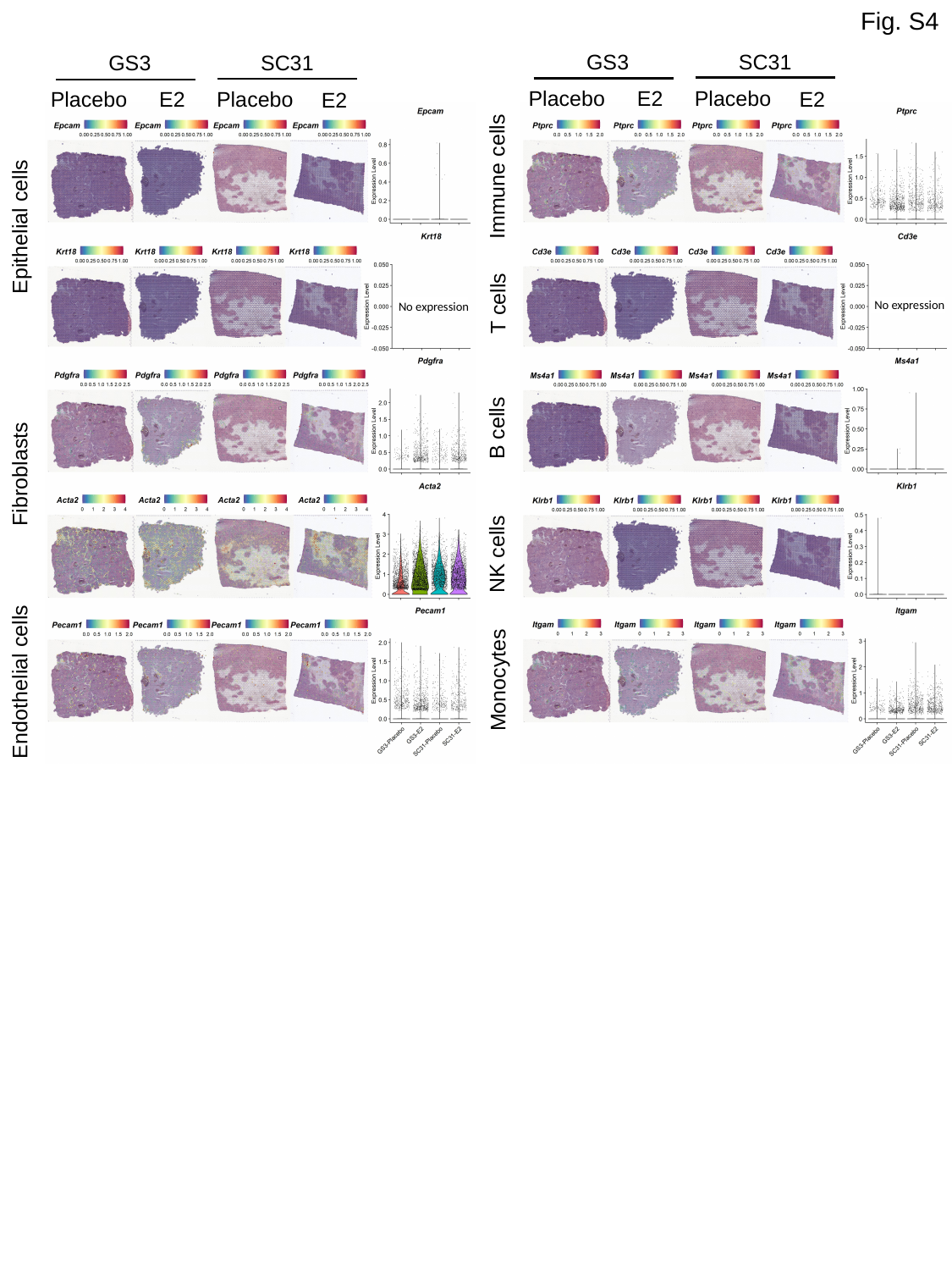

Fig. S4
GS3
SC31
GS3
SC31
E2
Placebo
Placebo
E2
E2
Placebo
Placebo
E2
Immune cells
Epithelial cells
T cells
No expression
No expression
B cells
Fibroblasts
NK cells
Monocytes
Endothelial cells

### Slide 5
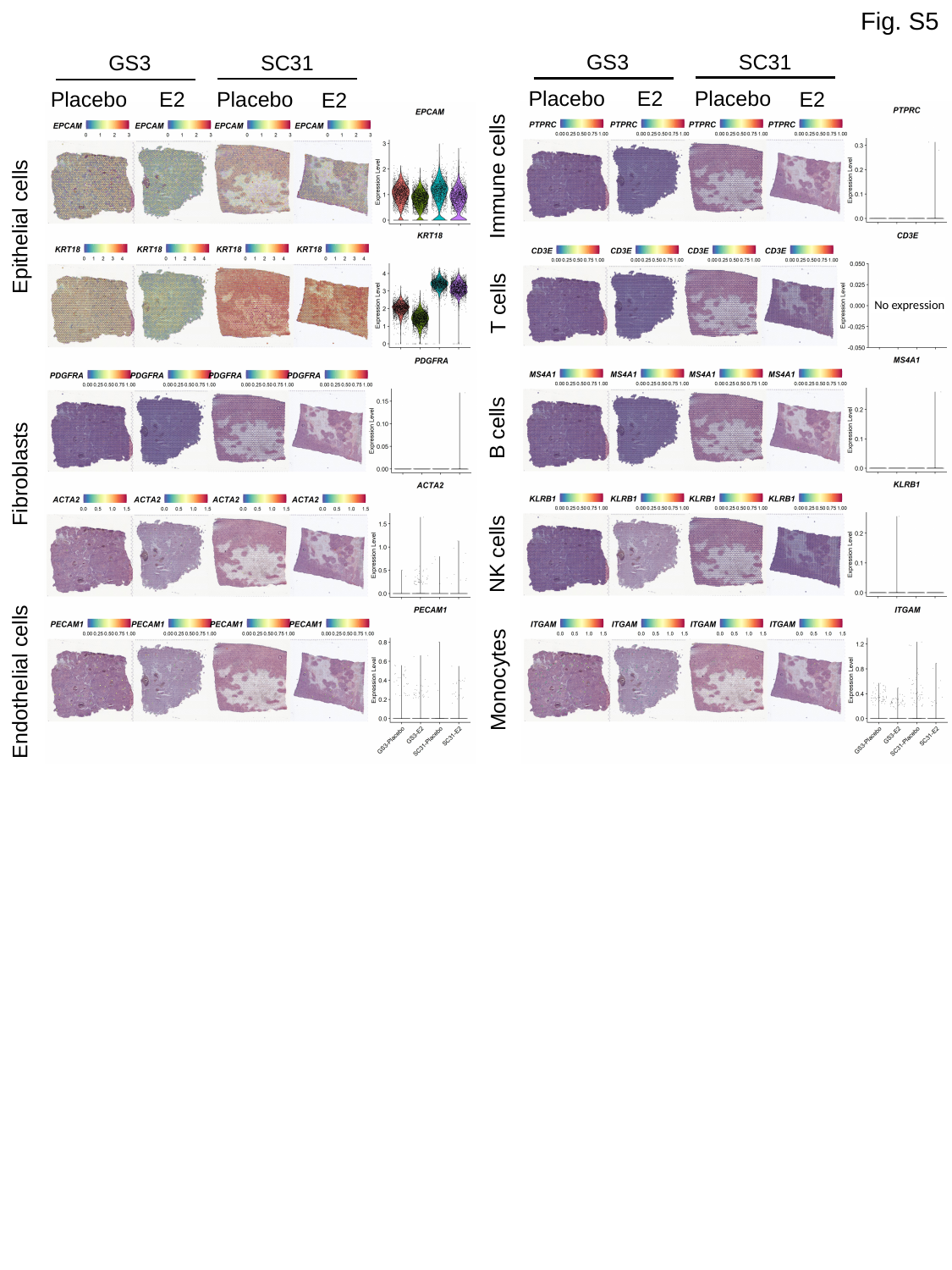

Fig. S5
GS3
SC31
GS3
SC31
E2
Placebo
Placebo
E2
E2
Placebo
Placebo
E2
Immune cells
Epithelial cells
T cells
No expression
B cells
Fibroblasts
NK cells
Monocytes
Endothelial cells

### Slide 6
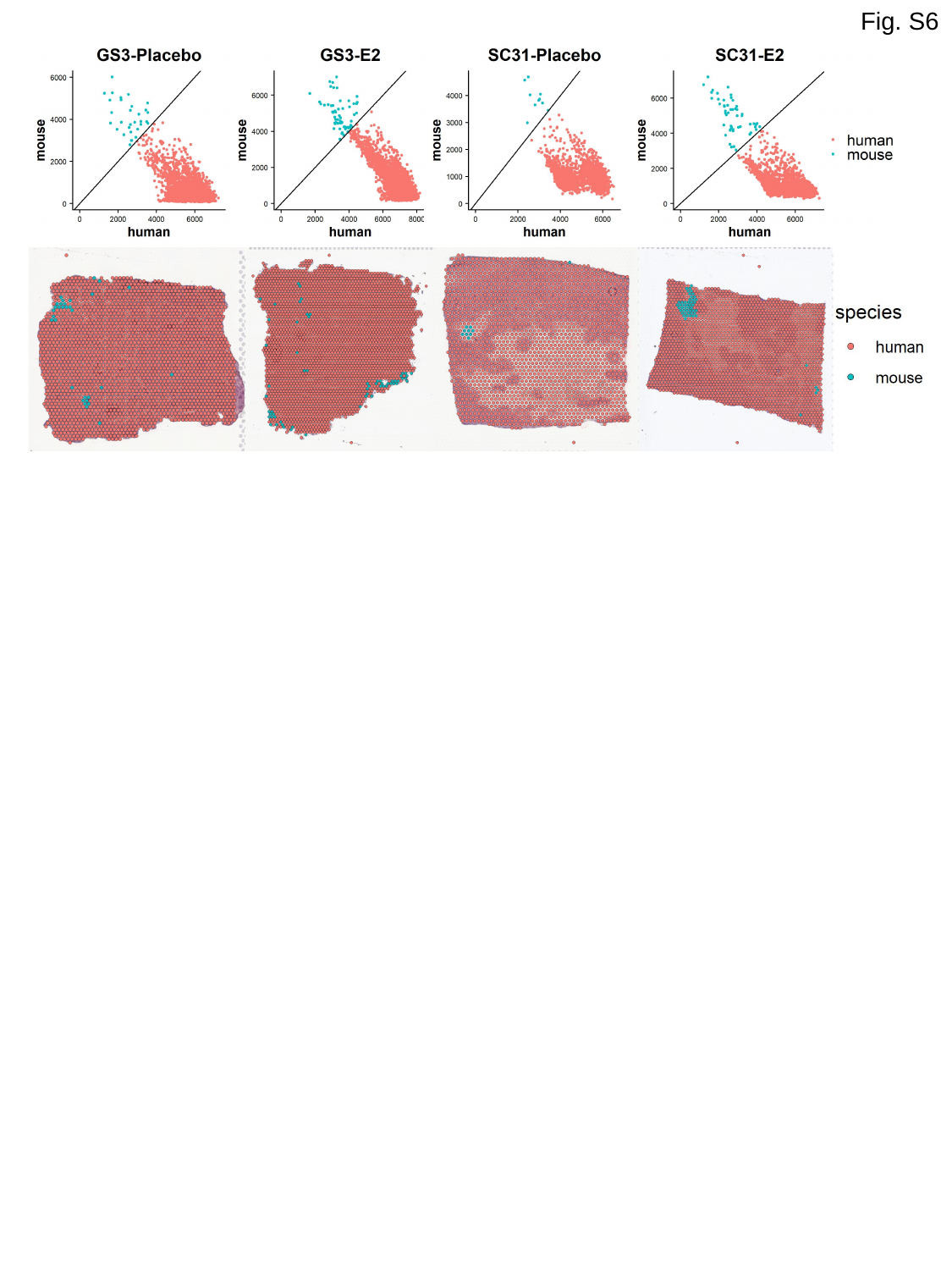

Fig. S6

### Slide 7
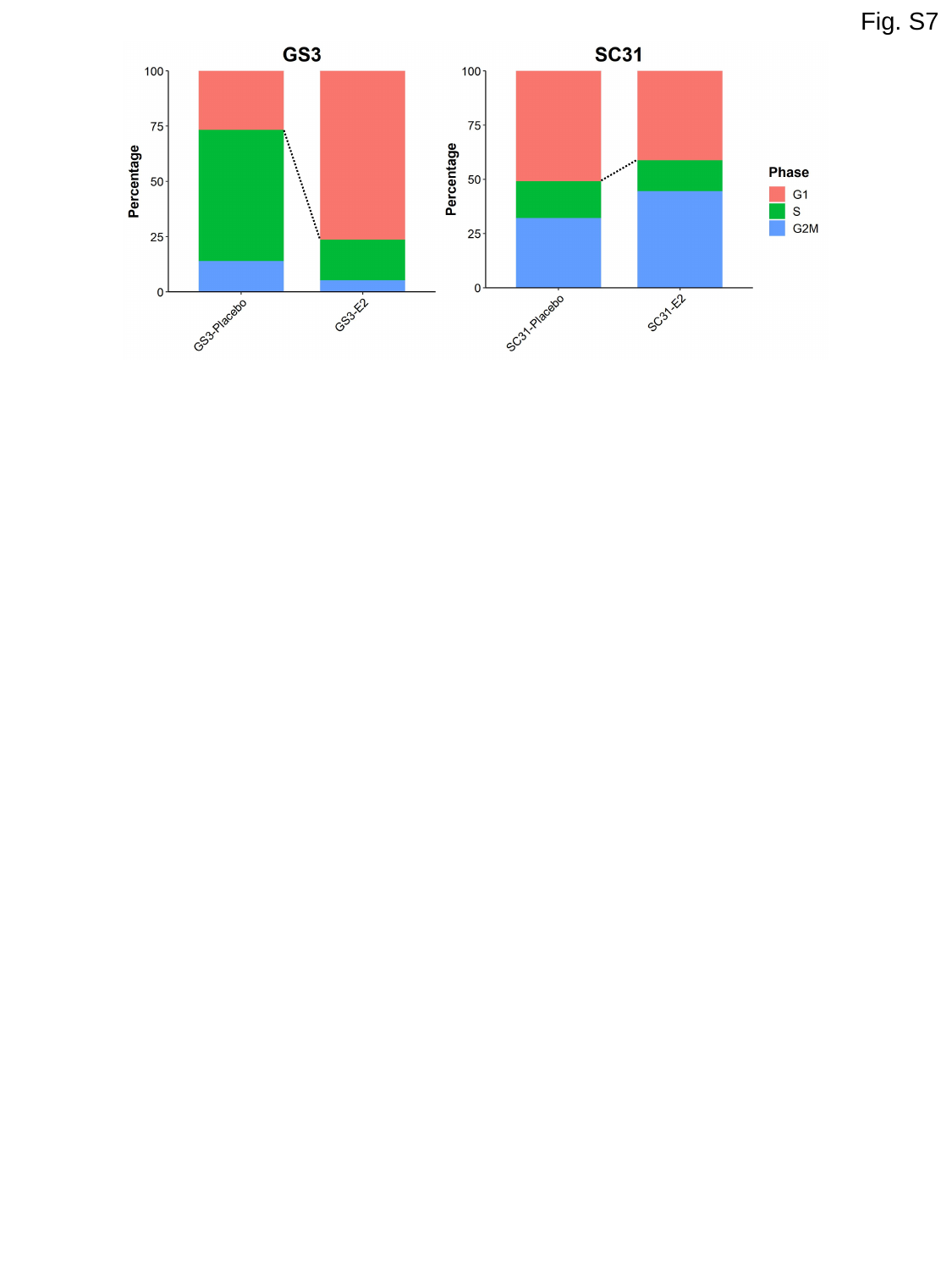

Fig. S7

### Slide 8
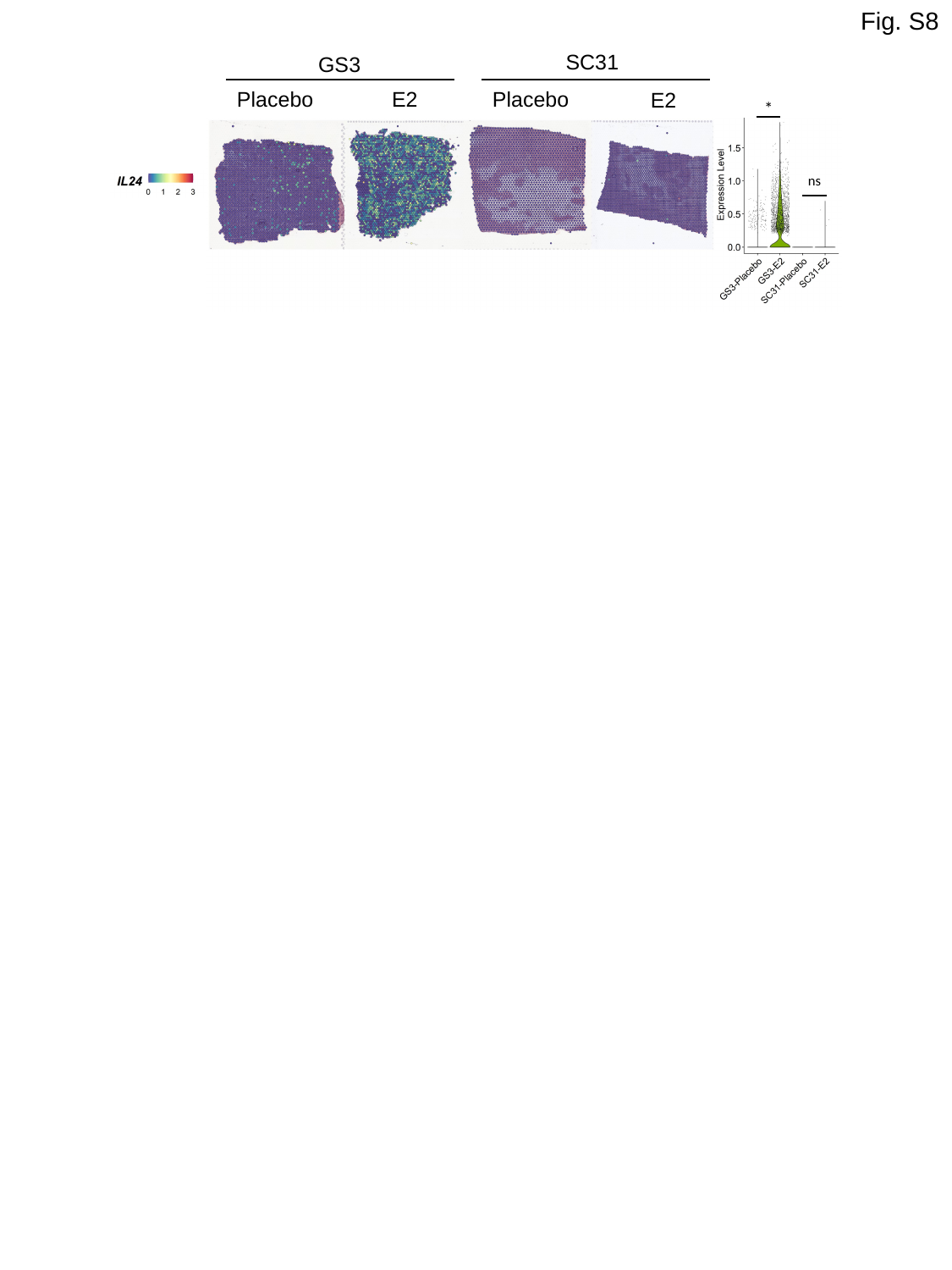

Fig. S8
SC31
GS3
E2
Placebo
Placebo
E2
*
ns

### Slide 9
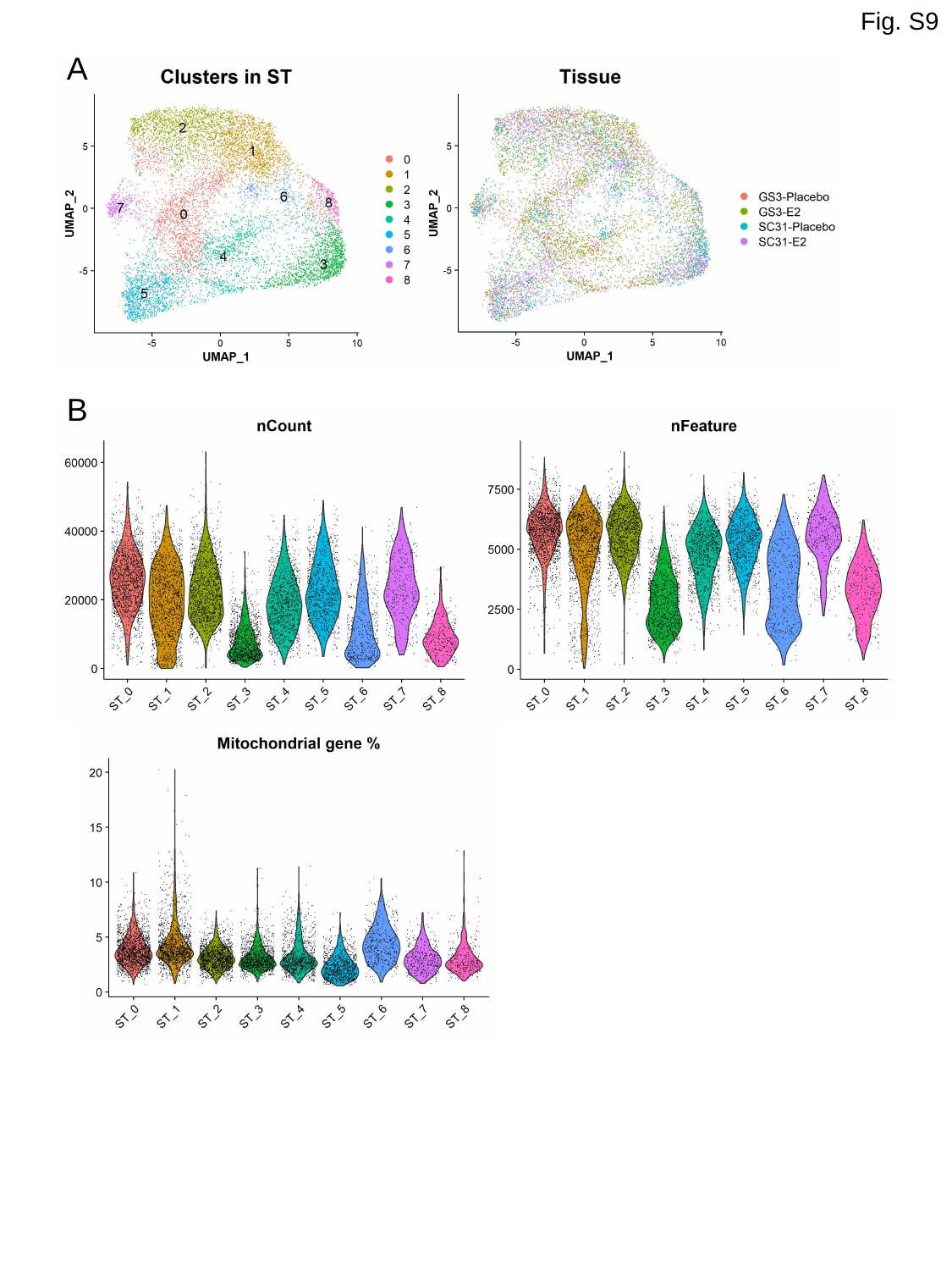

Fig. S9
A
B

### Slide 10
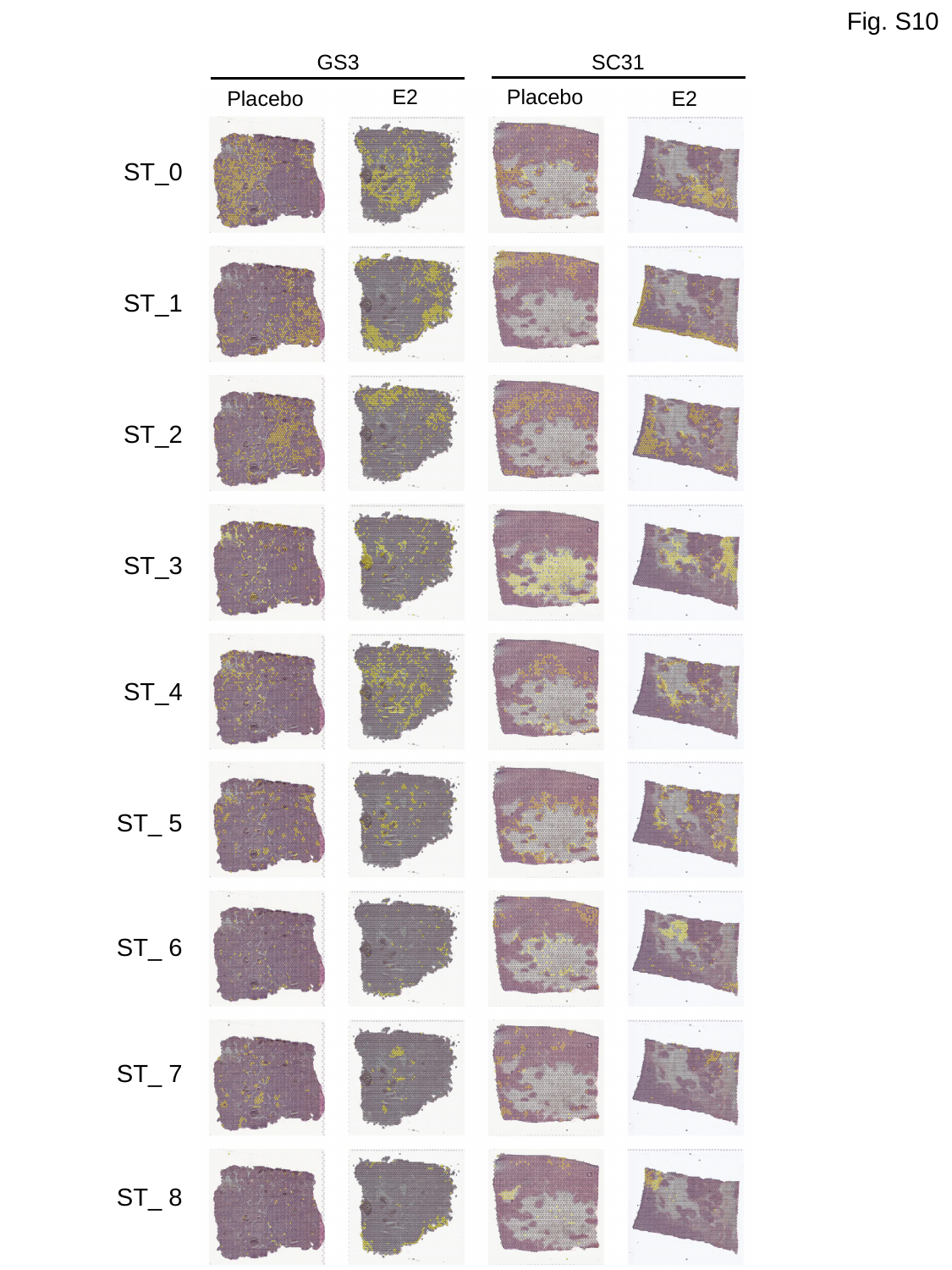

Fig. S10
GS3
SC31
E2
Placebo
Placebo
E2
ST_0
ST_1
ST_2
ST_3
ST_4
ST_ 5
ST_ 6
ST_ 7
ST_ 8

### Slide 11
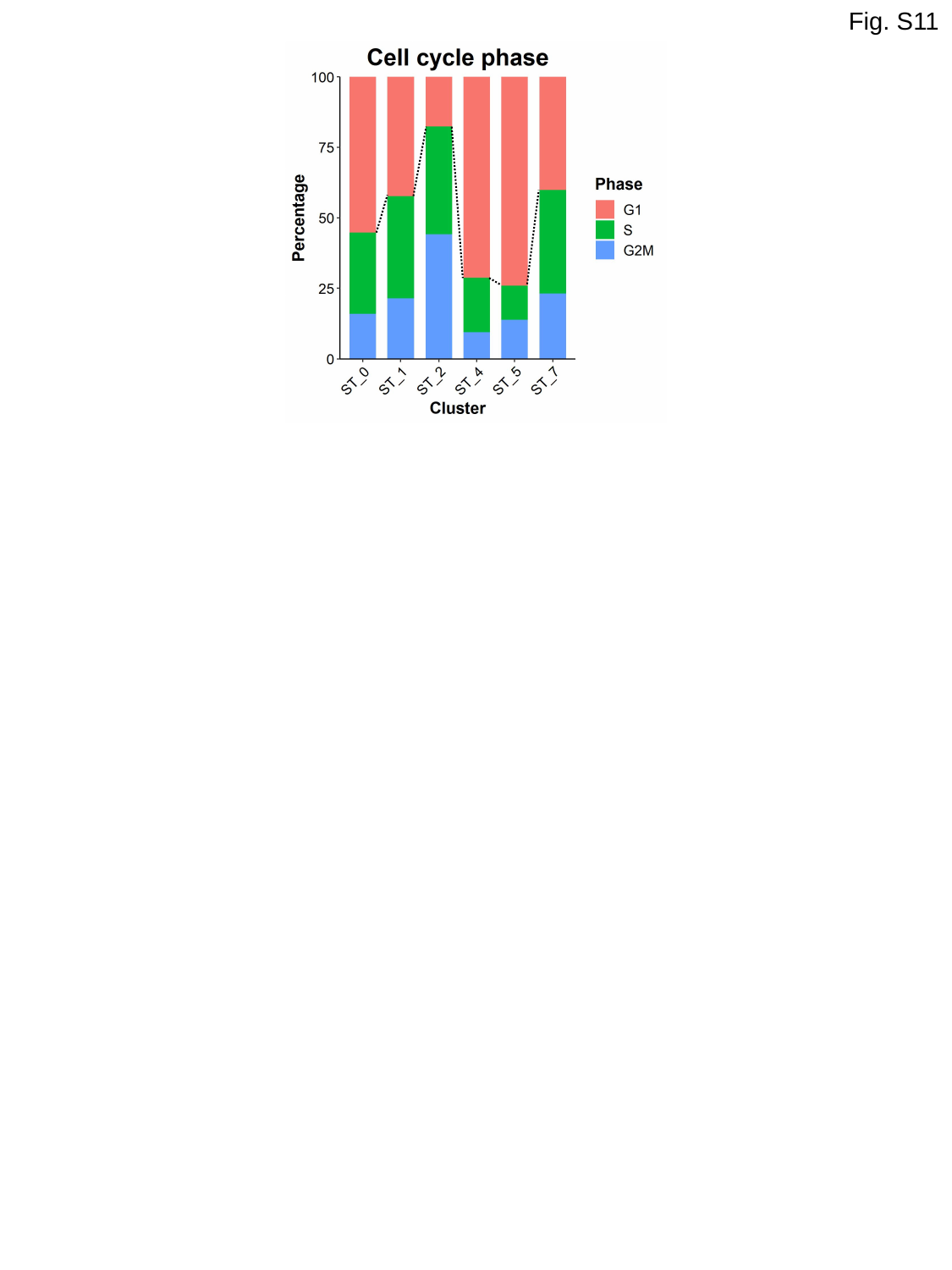

Fig. S11

### Slide 12
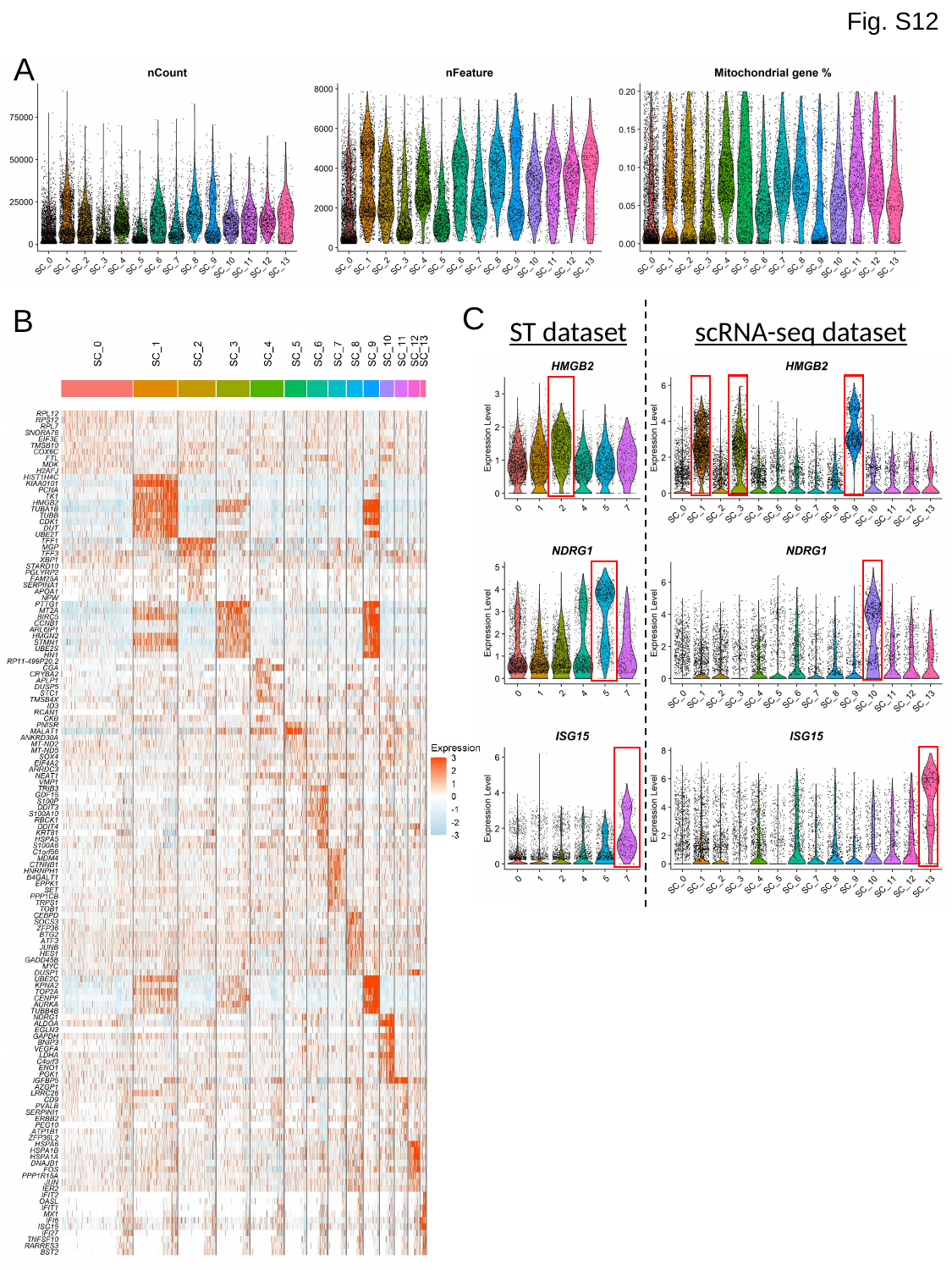

Fig. S12
A
C
B
ST dataset
scRNA-seq dataset

### Slide 13
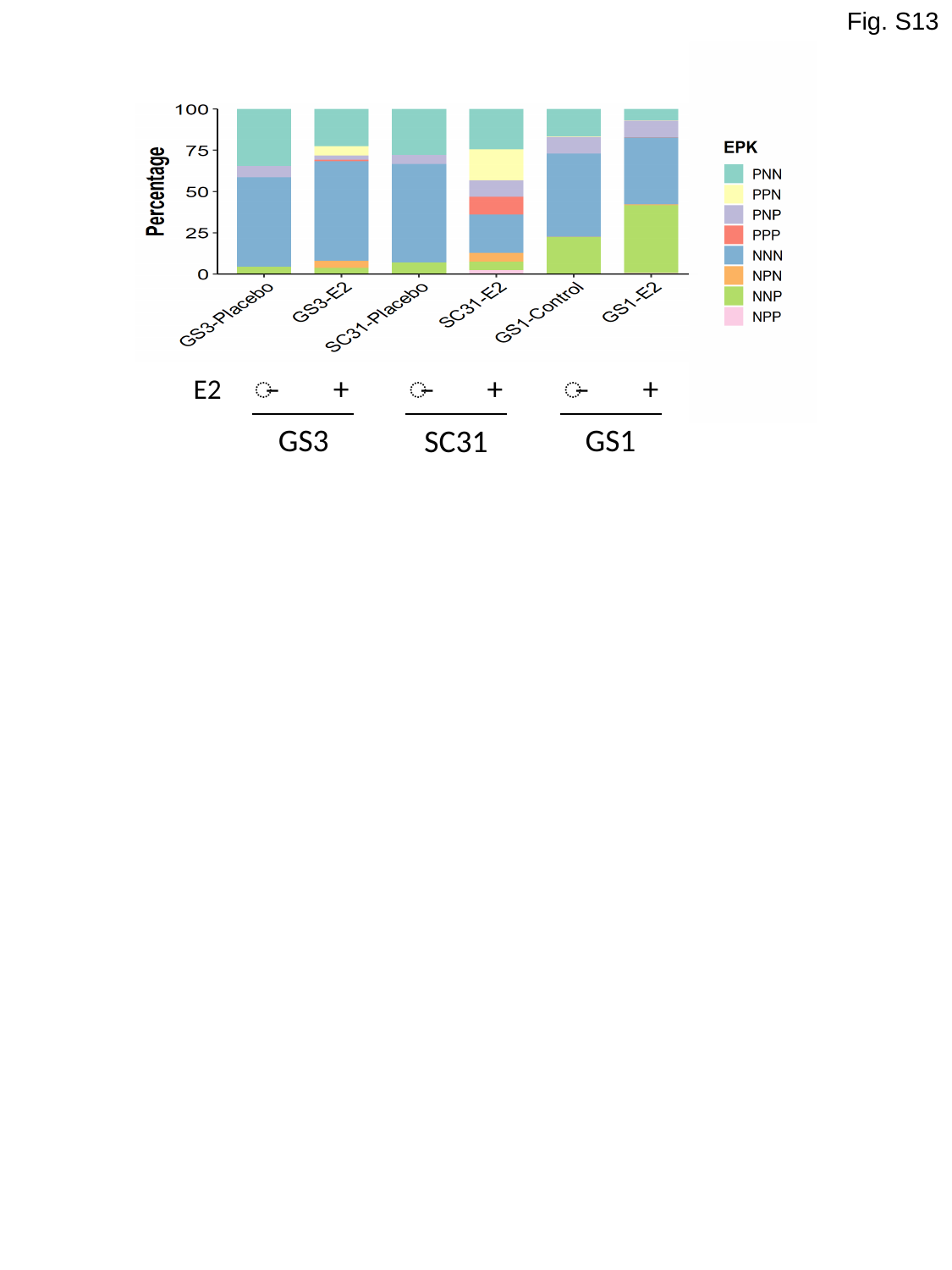

Fig. S13
̶
+
̶
+
̶
+
E2
GS3
GS1
SC31

### Slide 14
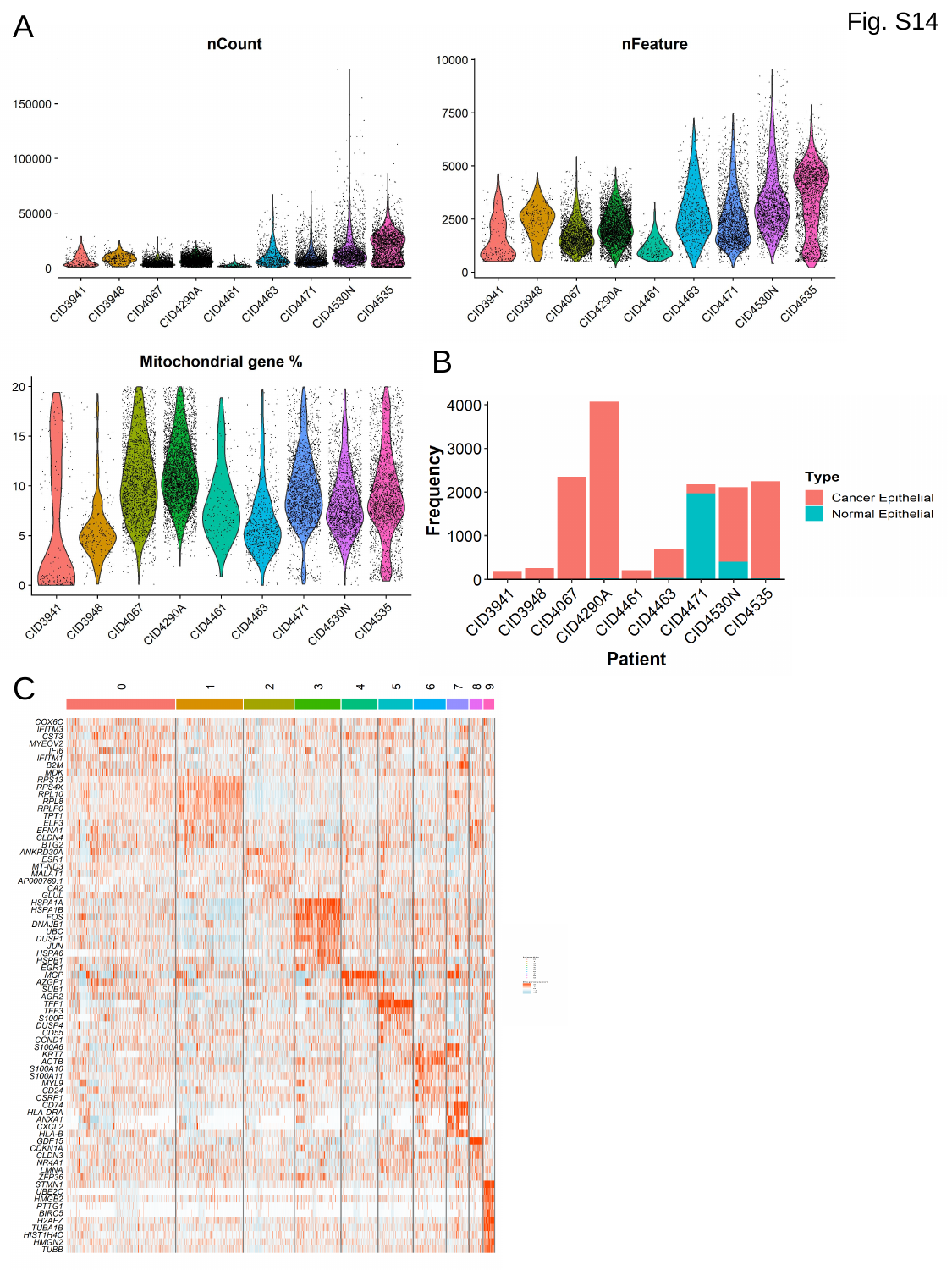

A
Fig. S14
B
C

### Slide 15
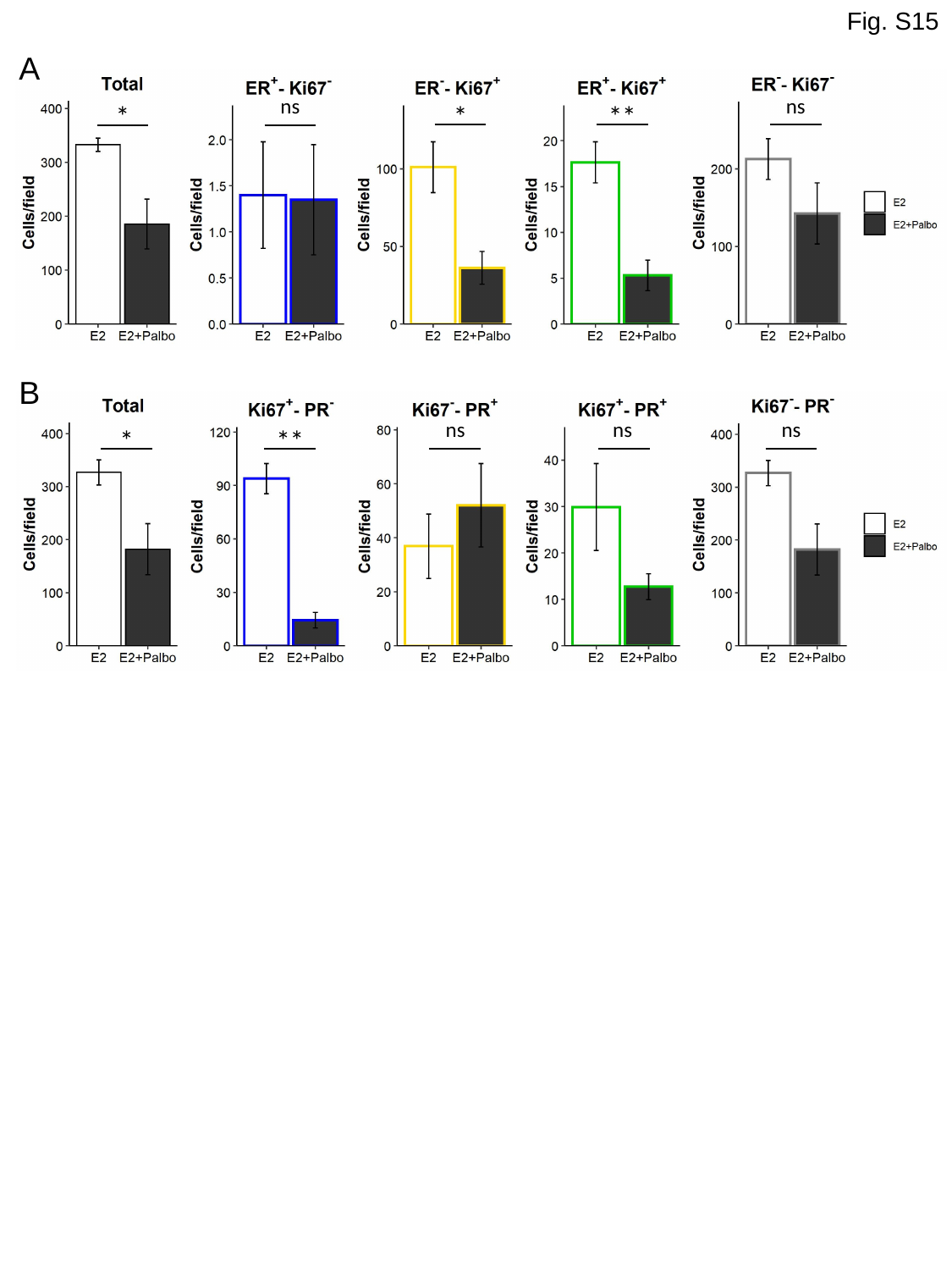

Fig. S15
A
ns
ns
*
*
**
B
ns
ns
ns
*
**

### Slide 16
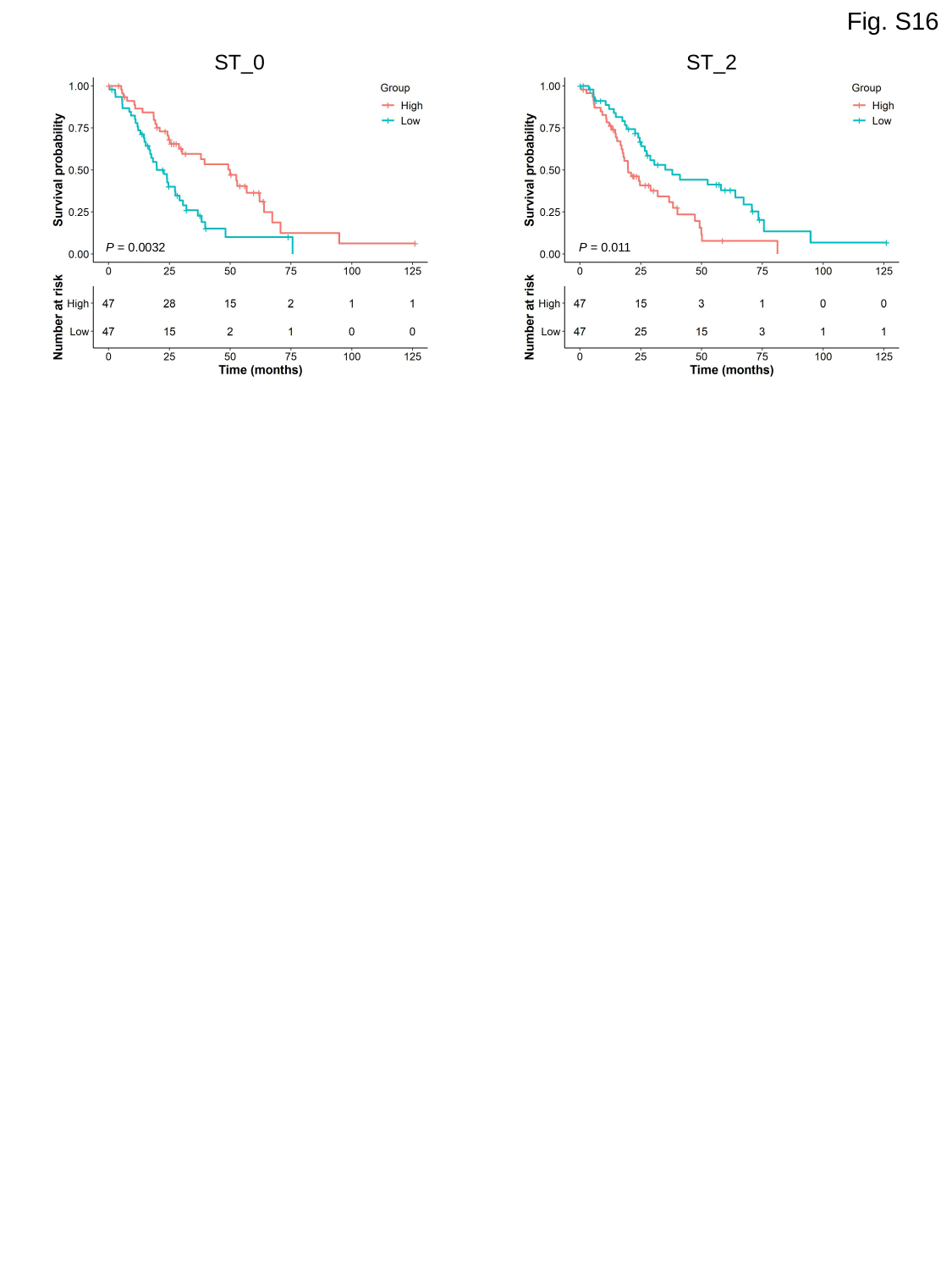

Fig. S16
ST_0
P = 0.0032
ST_2
P = 0.011

### Slide 17
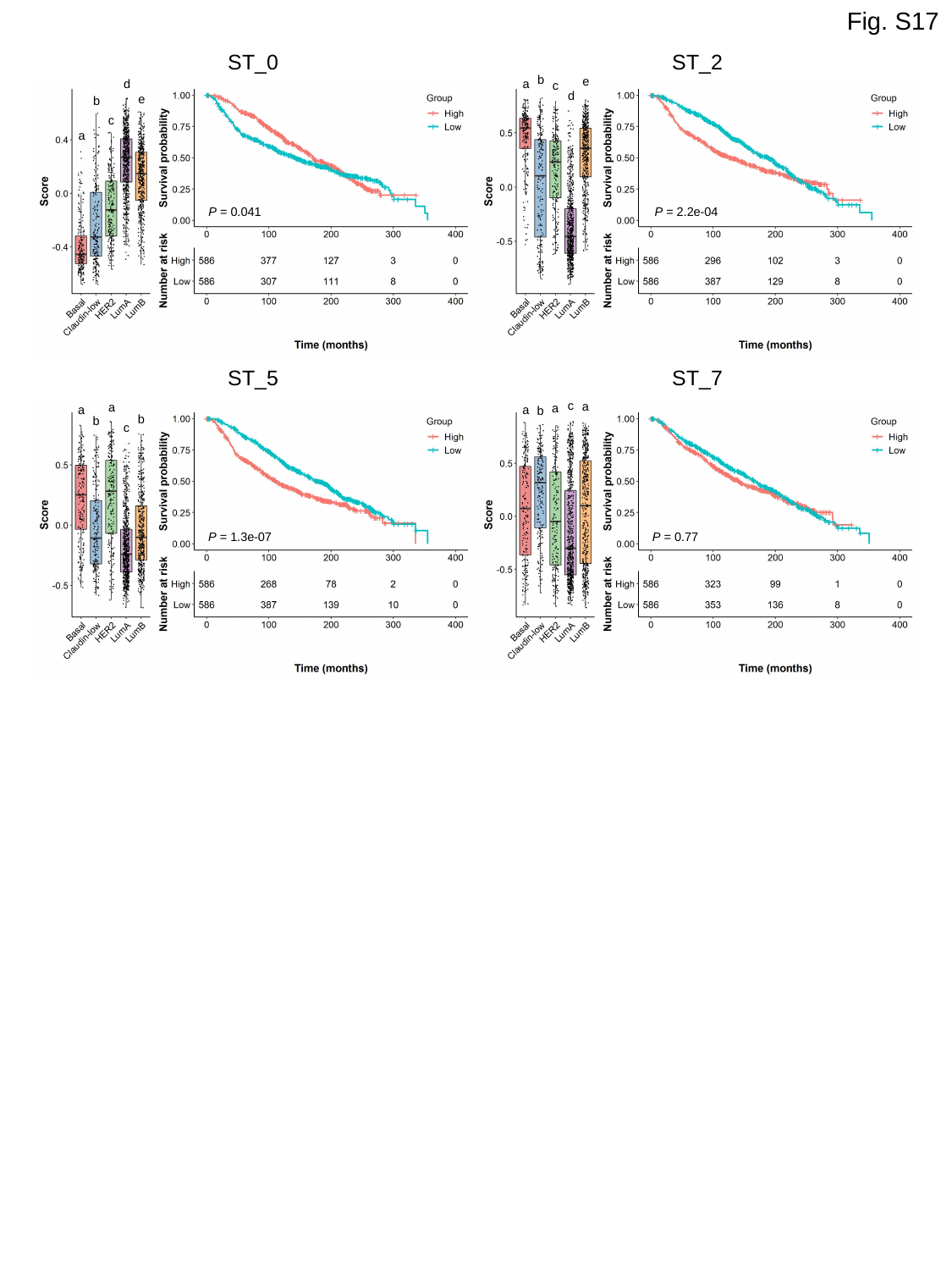

Fig. S17
ST_0
ST_2
b
e
d
a
c
d
e
b
c
a
P = 0.041
P = 2.2e-04
ST_5
ST_7
c
a
a
a
a
a
b
b
b
c
P = 1.3e-07
P = 0.77

### Slide 18
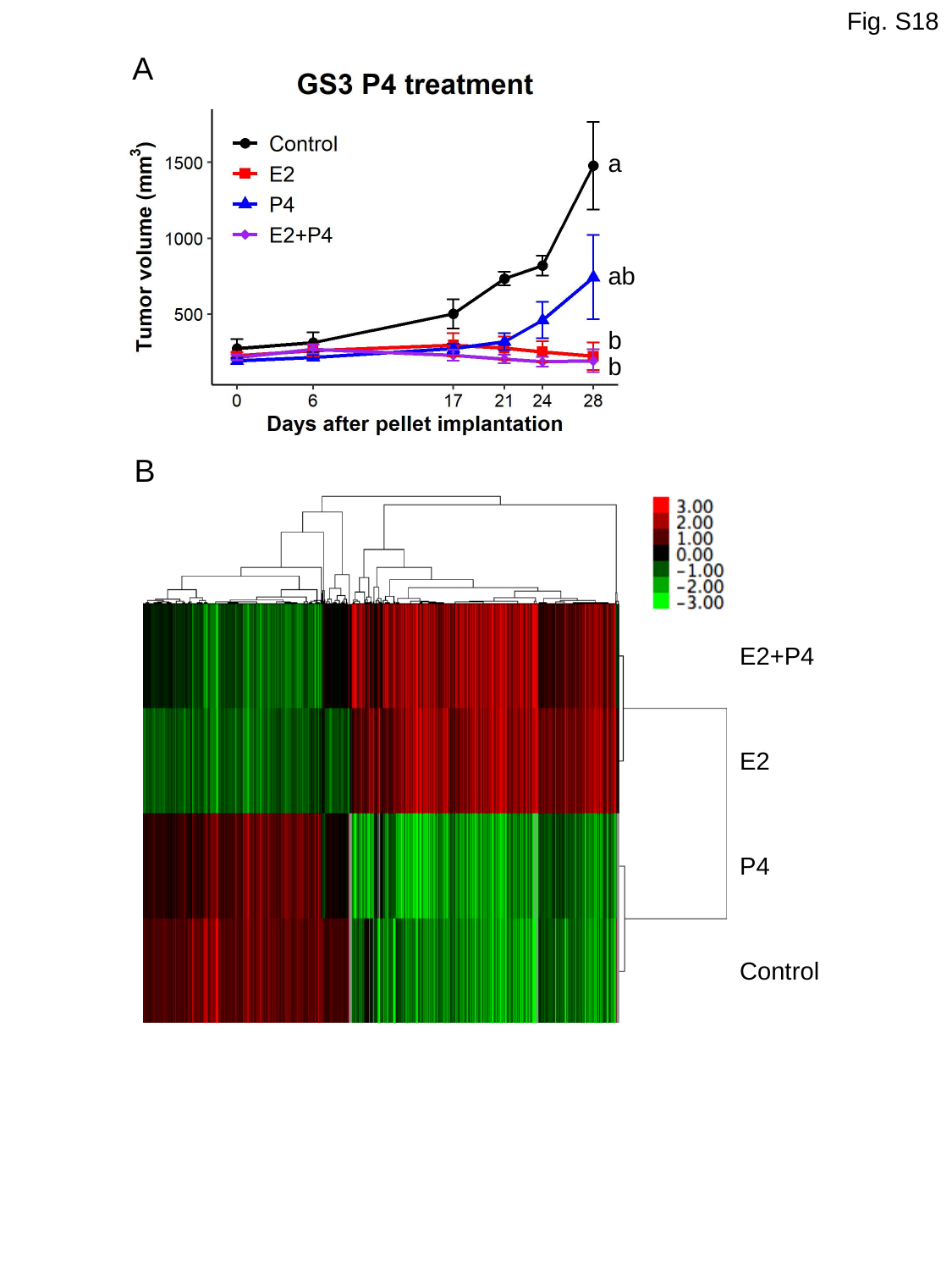

Fig. S18
A
a
ab
b
b
B
E2+P4
E2
P4
Control
