## Supplementary Figure legends for "Identification and characterization of a proliferative cell population in estrogen receptor-positive metastatic breast cancer through spatial and single-cell transcriptomics"

**Figure S1. Growth responses of ER^+^ breast cancer PDX models to estrogen treatment.**

**A** Growth curves of the E2-accelerating (GS1, GS2) and -dependent (GS4) breast cancer PDXs. Data are shown as mean ± SEM. *, *P* < 0.05; **, *P* < 0.01. **B** Tumor volume changes of GS3 and SC31 used for the ST experiment.

**Figure S2. Hematoxylin and eosin stained images of the sections used for the ST experiment.**

Scale bar = 2 mm.

**Figure S3. QC metrics of the datasets obtained from the ST experiment.**

The spatial plot on the left shows the location and level of the QC metric values on each section. The violin plot on the right summarizes the QC metrics in each section.

**Figure S4. Expression of mouse-derived stromal cell markers in the ST datasets.**

The spatial plot on the left shows the location and level of the expression on each section. The violin plot on the right summarizes the gene expressions in each section.

**Figure S5. Expression of human-derived stromal cell markers in the ST datasets.**

The spatial plot on the left shows the location and level of the expression on each section. The violin plot on the right summarizes the gene expressions in each section.

**Figure S6. Comparison of human- and mouse-derived gene expression in the ST datasets.**

The proportion of human- and mouse-derived genes in each spot. Red and blue spot represents the spot including more human and mouse genes, respectively.

**Figure S7. Cell cycle phases of the spots in the ST datasets.**

**Figure S8. Expression of *IL24* in the ST datasets.**

The spatial plot on the left shows the location and level of the expression on each section. The violin plot on the right summarizes the gene expressions in each section. *, *P* < 0.05; ns, not significant.

**Figure S9. Clustering on the integrated ST dataset.**

**A** UMAP plot of the integrated dataset. Spot is represented by the dot and is colored according to the cluster (ST_0–8) identified by the unbiased clustering (left) or the originated datasets (right). **B** QC metrics of the ST clusters.

**Figure S10. Localization of the ST clusters on each section.**

Yellow dot shows the localization of the ST cluster (ST_0-8) indicated on the left.

**Figure S11. Cell cycle phases of the spots included in each ST cluster.**

**Figure S12. Analyses on the integrated scRNA-seq dataset from GS3 and SC31.**

**A** QC metrics of the clusters (SC_0-13) identified in the scRNA-seq dataset. **B** Heatmap of the top 10 genes for the SC clusters. Column represents the cells and row represents the genes. The gene expression levels were scaled by SCTransform function. **C** Comparison of expression patterns of the cluster-specific genes between the ST and scRNA-seq datasets.

**Figure S13. EPK classification on the GS3/SC31/GS1 scRNA-seq datasets.**

**Figure S14. Analyses on the integrated scRNA-seq dataset from human ER^+^ breast cancers.**

**A** QC metrics of the individual datasets in the integrated human scRNA-seq dataset (n = 9). **B** The number of normal or cancer epithelial cells from the individual datasets. **C** Heatmap of the top 10 genes for the clusters. Column represents the cells and row represents the genes. The gene expression levels were scaled by SCTransform function.

**Figure S15. Quantification of the dual IHC for ER/Ki67 and Ki67/PR.**

The dual IHC results for **A** ER/Ki67 and **B** Ki67/PR. Bar plot indicates the number of the cells classified as indicated per field. Data are shown as mean ± SEM [n = 4 (E2) and 5 (E2 + Palbo)]. *, *P* < 0.05; **, *P* < 0.01; ns, not significant.

**Figure S16. Clinical data analysis with ST_0 and ST_2 signatures on GSE124647 cohort.**

Stage IV ER^+^/HER2^-^ breast cancers in the GSE124647 cohort (n = 140) were analyzed using ST_0 and ST_2 signatures. The gene signature scores were calculated in each patient using GSVA R package. Kaplan-Meier plots show the overall survival of patients in each group.

**Figure S17. Clinical data analysis with the ST signatures on all breast cancer subtypes in METABRIC cohort.**

All breast cancers in the METABRIC cohort (basal-like, n = 199; claudin-low, n = 199; HER2-enriched, n = 220; luminal A, n = 679; luminal B, n = 461) were analyzed using the gene signatures from ST_0, ST_2, ST_5, and ST_7. The gene signature scores were calculated in each patient using GSVA R package. Left panels show box plots of the scores in each subtype. Different letters on the box plots indicates significant difference between the subtypes (*P* < 0.05). Right panels show Kaplan-Meier plots for the overall survival of the patients in each group.

**Figure S18. Effect of progesterone treatment on GS3 model.**

**A** Growth curve of GS3 in control, E2, P4, and E2 + P4 groups. Data are shown as mean ± SEM. Different letters indicates significant difference between the groups on day 28 (*P* < 0.05). **B** Heatmap of the results from bulk RNA-seq analysis of GS3 in control, E2, P4, and E2 + P4 groups. Row represents the samples and column represents the genes. The expression levels of the genes were scaled per column.
